# EP300 (p300) mediated histone butyrylation is critical for adipogenesis

**DOI:** 10.1101/2021.08.01.454641

**Authors:** Aditya Bhattacharya, Sourav Chatterjee, Utsa Bhaduri, Akash Kumar Singh, Madavan Vasudevan, Koneni V Sashidhara, Rajdeep Guha, Aamir Nazir, Srikanta Kumar Rath, Nagashayana Natesh, Tapas K Kundu

## Abstract

**Objective:** The master epigenetic enzyme EP300 (p300) besides having lysine acetyltransferase activity can also catalyse other acylation modifications (propionylation, butyrylation, crotonylation etc.), the physiological implications of which are yet to be established fully. We hypothesized that p300 catalysed histone butyrylation may have a causal relationship with adipogenesis and the consequent obesity.

**Methods:** Histone butyrylation pattern was investigated in 3T3L1 cells upon adipogenesis by immunoblotting and chromatin immunoprecipitation experiments. A small molecule modulator that could specifically inhibit p300 catalysed butyrylation without affecting its canonical acetyltransferase activity was screened from a series of compounds and then administered in differentiating 3T3L1 adipocytes as well as high fat diet-induced and genetically obese mice to validate the importance of butyrylation in adipogenesis.

**Results:** Histone butyrylation was increased upon adipogenesis both globally and locally in the promoters of pro-adipogenic genes along with an upregulation in the expression of acyl CoA generating enzyme Acss2, knockdown of which led to reduced butyrylation. Treatment of differentiating 3T3L1 cells with the p300 specific butyrylation inhibitor LTK-14A led to abrogation of adipogenesis with reduced expression of pro-adipogenic genes and inhibition of H4K5 butyrylation. LTK-14A administration could also attenuate weight gain in both mice models of obesity by preventing adipocyte hypertrophy via H4K5 butyrylation inhibition.

**Conclusion:** Our results indicate that p300 catalysed histone butyrylation may have a causal relationship with the process of adipogenesis. Site specific inhibition of butyrylation could lead to adipogenesis repression and hence this epigenetic modification could be targeted for obesity treatment.

**Highlights:**

- Histone butyrylation has been established as a new epigenetic signature in the context of adipogenesis.
- To the best of our knowledge, this is the first report of a selective inhibitor of p300 catalysed histone acylation (butyrylation) without affecting its canonical acetyltransferase activity.
- Specific inhibition of H4K5 butyrylation could be a possible mechanism for inhibiting adipogenesis and hepatic steatosis leading to better control of obesity.
- LTK-14A class of molecule could be developed as anti-obesity therapeutics.

**Graphical abstract:** Proposed model for the role of p300-mediated histone butyrylation in adipogenesis:
In pre-adipocytes, there exists a basal level of histone acetylation while butyrylation is present to a much lesser extent owing to low stoichiometric levels of butyryl CoA. Induction of adipogenesis causes a simultaneous upregulation of histone acetylation and butyrylation marks leading to increased rate of adipogenesis and concomitant transcriptional activation of pro-adipogeneic genes.
Onset of obesity in mice, either due to excess energy intake through high fat diet consumption or increased de novo synthesis of fatty acids due to leptin receptor gene mutation leading to hyperphagic behavior, is accompanied by adipocyte hyperplasia and hypertrophy. Both the organs of adipose tissue and liver were found to have enhanced levels of H4K5 butyrylation during obesity. LTK-14A, a butyrylation specific inhibitor could efficiently prevent the processes of adipogenesis and adipocyte hypertrophy due to inhibition of H4K5 butyrylation in these organs. Thus the compound could attenuate weight gain by selective inhibition of butyrylation without affecting acetylation, thereby highlighting the importance of histone butyrylation in adipogenesis and obesity.

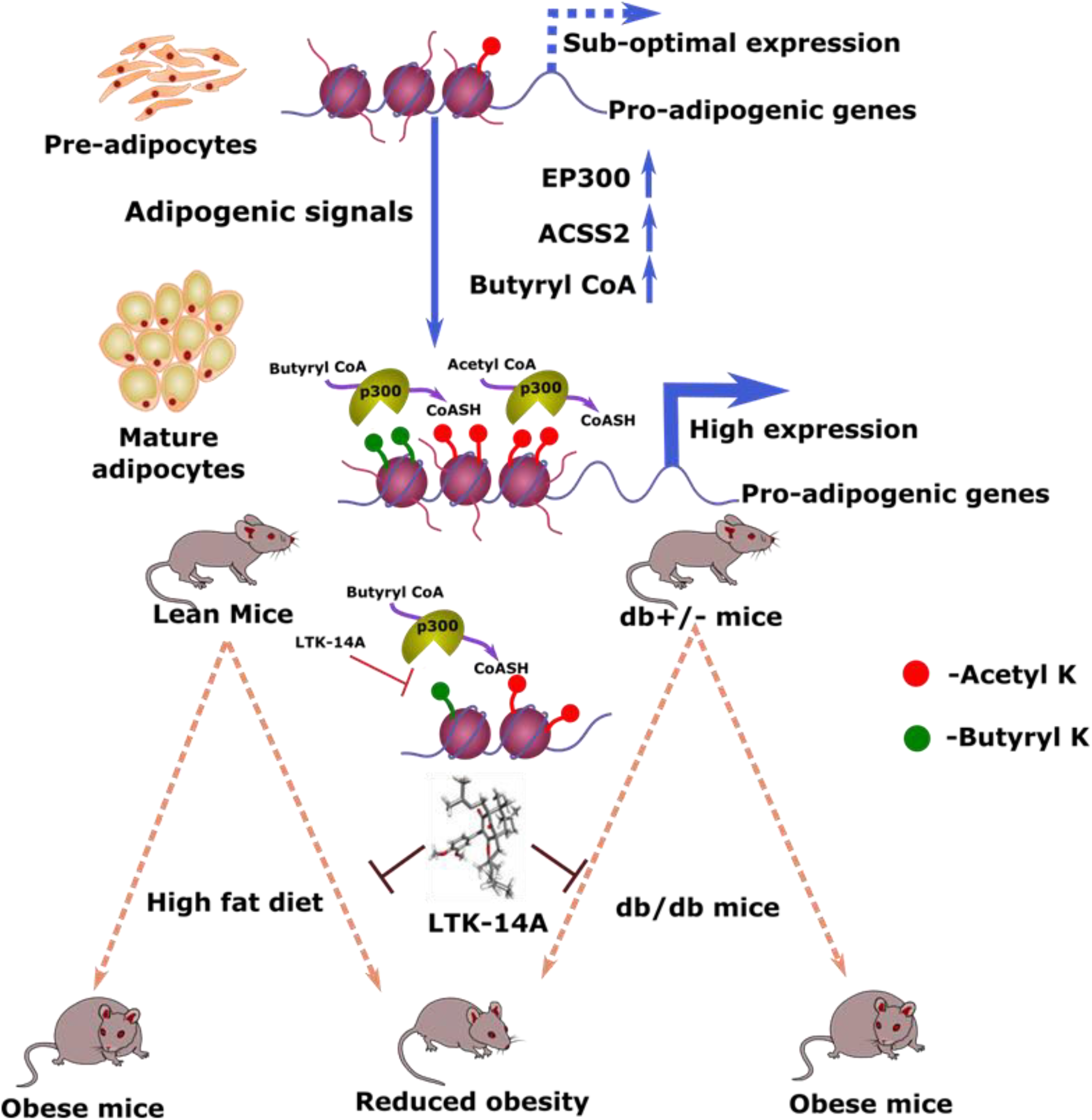

## Introduction

Chromatin function is spatially and temporally regulated by several processes like incorporation of histone variants, remodeling of nucleosomes by ATP-dependent remodelers and histone post-translational modifications. Interestingly, epigenetics and the metabolic state of the cell are intimately intertwined because the enzymes catalyzing the modifications use co-factors derived from several metabolic reactions (1). In recent times, advancements made in high sensitivity mass spectrometry have led to the discovery of new acylation modifications that structurally resemble acetylation, but differ in hydrocarbon chain length, hydrophobicity and charge (2). Emerging evidence indicates that these modifications occur dynamically on histones and could either have distinct functions from acetylation, or, have a similar but more profound impact on chromatin de-compaction, thereby amplifying the transcriptional output (3). EP300 (p300) is the master acyltransferase that has been found to promiscuously acylate histones using diverse CoA derivatives (2).

Due to the rising prevalence of obesity in the modern-day society, there has been increased research on the molecular mechanisms behind its manifestation. At the core of obesity lies the adipose tissue which regulates energy homeostasis and metabolism. Hence, the developmental process of adipose tissue i.e. adipogenesis and its dysregulation needs to be studied for clinical management of obesity-related disorders. The process of adipogenesis is accompanied by a significant increase in global histone acetylation levels, along with a rise in expression of p300 and a concomitant decrease in the levels of several histone deacetylases (4). Regulation of gene expression is dictated by the fluctuating activities of histone acetyltransferases and histone deacetylases. Moreover, the importance of histone acetylation for adipogenesis has been further underlined by the finding that knockdown of ATP citrate lyase, an enzyme responsible for the regeneration of acetyl CoA in the cytoplasm and nucleus, leads to decrease in histone acetylation and also suppression of adipogenesis (5). Therefore, it can be assumed that p300 plays a crucial role in promoting the expression of adipogenesis related genes by acetylating the histones in the corresponding gene regulatory regions, especially the promoters. Amongst the different coenzyme A derivatives, butyric acid is an important short chain fatty acid and an intermediate of fatty acid synthesis/breakdown processes. p300 has been known to carry out butyrylation of histones which has been causally linked to the process of spermatogenesis (3).

In this study we report that histone butyrylation is critical in adipoogenesis, probed by the first known acylation-specific small molecule inhibitor of p300. The newly discovered importance of histone butyrylation in adipogenesis makes it an attractive target for small molecule compound mediated inhibition of adipogenesis, thereby controlling obesity with lesser collateral damage.

## Materials and Methods

### Animals and treatment for obesity-related studies

Male C57BL/6J mice (8 weeks old) were bred in - house and acclimatized for 1 week prior to experiments. Mice were maintained on a 12 hour light/dark cycle and given ad libitum access to food and water. Experiments were carried out in accordance with the Committee for the Purpose of Control and Supervision of Experiments on Animals (CPCSEA) guidelines and were approved by the Institutional Animal Ethics Committee of JNCASR (201/GO/Re/S/2000/CPCSEA dated 1st June, 2000 Renewed in Aug 2015). The age-matched animals were maintained on a standard laboratory chow diet (AMRUT M/s Krishna Valley Agrotech, Maharashtra) or high fat diet (60 kcal% fat) (Research Diets, Cat. No. D12492) with or without LTK-14A (50 mg/Kg body weight) and water. The compound was mixed with the high fat diet according to dosage requirements in the laboratory every week. Body weight and food intake were measured twice every week.

For the experiment on genetically obese mice, male db/db mice and db+/- mice (8-10 weeks old) bred in-house were obtained and acclimatized for 1 week prior to experiments. Mice were maintained on a 12-hour light/dark cycle at 22+/-3 ᵒC and a relative humidity of 55% and given ad libitum access to food and water. The experiments were carried out in accordance with the Committee for the Purpose of Control and Supervision of Experiments on Animals (CPCSEA) guidelines, Government of India and were approved by the Institutional Animal Ethics Committee of CSIR-CDRI, Lucknow (IAEC/2020/115/Renew-0/Sl.No.09). The age-matched animals were maintained on a standard laboratory pelleted diet (Altromin International, Cat. No. 1320) and water. db/db mice were oral gavaged (50 mg/kg) with LTK14A in 0.5% carboxymethyl cellulose, an equal volume of the vehicle was orally gavaged to control db/db mice for a total period of 30 days. db+/- animals were kept as an untreated normal control group. The body weights and feed intake was recorded every week throughout the experimental period.

### Cell culture

The mammalian cell line 3T3L1 was obtained from ATCC (American Type Culture Collection) and was grown in DMEM (Dulbecco’s Modified Eagle’s Medium). The growth medium was supplemented with 10% FBS (Fetal Bovine Serum). The cell line was grown in 37°C incubator with 5% CO2 and 90% relative humidity.

### Adipogenesis and Oil Red-O staining

3T3L1 preadipocytes were grown to confluency and 24 hours post-confluence, cells were incubated with a differentiation medium (growth medium containing 1µM Dexamethasone, 0.5 mM IBMX and 1 µg/ml insulin)(Sigma Aldrich) for 2 days in the presence/absence of LTK-14A. Then the cells were supplemented with a maintenance medium with/without LTK-14A (growth medium with 1 µg/ml insulin only) every alternate day. On the 6th day post-induction of differentiation the cells were fixed with 3.7% formaldehyde solution followed by staining with Oil Red-O dye (Sigma-Aldrich) (prepared in 60% isopropanol) for 1 hour at room temperature. The incorporated dye was extracted using isopropanol and absorbance values were determined spectrophotometrically at 510 nm to estimate total amount of lipid accumulation under different experimental conditions.

### Acid extraction of histones and immunoblotting

3T3L1 cells were washed with ice-cold PBS and then resuspended in Triton extraction buffer (PBS containing 0.5% Triton X-100 (v/v) and 2 mM PMSF). The cells were allowed to lyse on ice for 10 minutes, spun down, washed with PBS and the process was repeated another time. Then the proteins were acid extracted with 0.2 N HCl for two hours on ice. The debris was spun down and then histones were precipitated by incubating the supernatant with 33% trichloroacetic acid (Sigma-Aldrich) at 4ᴼC for 30 minutes. The histones were then pelleted down and then given two washes with acetone. The pellet was allowed to air-dry for 10 minutes and then resuspended with 50 mM Tris-Cl, pH 7.4. Extraction of histones from epididymal fat pads was performed by mechanically homogenizing the adipose tissues in Triton extraction buffer and then following the same subsequent steps as mentioned before, except for the step of acid extraction with HCl, in which the incubation was performed for four hours on ice.

The samples were then run in 15% SDS-PAGE, transferred onto PVDF membrane (Milipore) in transfer buffer (25 mM Tris, 192 mM glycine, 0.036% SDS in 20% methanol), blocked with 5% skimmed milk solution to prevent non-specific antibody binding and finally probed with antibodies against acetylation and butyrylation marks prepared in 1% skimmed milk or FBS solution (5% BSA, 1x TBS, 0.1% Tween-20). HRP-conjugated secondary antibodies (Abcam) were used against the primary antibodies to obtain a chemiluminiscent signal in the presence of SuperSignal West Pico Chemiluminiscent Substrate (Thermo Scientific) which was developed using the VersaDoc imaging system (Biorad).

### Synthesis of LTK14A

Isolation of garcinol from Garcinia indica and synthesis of isogarcinol from it were carried out following the protocols as mentioned in Mantelingu et al, (2007) (11). For LTK-14A synthesis, anhydrous potassium carbonate (0.41 mmol) was added to a stirred solution of isogarcinol (100 mg, 0.16 mmol) in dry acetone. The mixture was stirred for 1 hour at room temperature followed by dropwise addition of dimethyl sulfate (0.41 mmol) under a nitrogen gas atmosphere. The resulting reaction mixture was further stirred at room temperature for 14 hours. Progress of the reaction was monitored through thin layer chromatography. Acetone was evaporated from the reaction mass, ice was added and acidified with 5% aqueous HCl (pH ∼ 5-6). A pale brown precipitate was observed, which was filtered and washed with cold water. The dry mass was then chromatographed on silica gel using 5-6% ethyl acetate in hexane as an eluting solvent mixture. 720 mg (69%) LTK14A was isolated as a white solid. NMR spectral analysis was consistent with the desired product (Sup. Fig.3A). A 20 mg sample was dissolved in hot methanol and placed in a flat surface glass vial with access to very slow solvent evaporation. Crystals as colorless needles were isolated. The purity of the synthesized LTK-14A was found to be more than 98% as verified by HPLC (Sup. Fig. 3C) and only the protonated species of LTK-14A were observed in the ESI spectra of the corresponding fraction (Sup. Fig. 3B).

### X-ray crystal structure determination for LTK14A (CCDC: 1969185)

Suitable single crystal with approximate dimensions of 0.35 × 0.18 × 0.15 mm3 was used for X-ray diffraction analyses by mounting on the tip of a glass fibre in air. Data were collected on a Bruker kappa apex 2 with Mo Kα (λ=0.71073 Å) at 296 (2) K. The structure was solved by direct method using program SHELXL-97 and subsequent Fast Fourier Transform technique. Crystallographic data and experimental details for LTK14A are summarized in Table 1 and Sup. Fig. 3D.

**Table 1:** Crystal data and structure refinement for LTK14A (CCDC: 1969185)

|  |  |
| --- | --- |
| Empirical formula | C<sub>40</sub>H<sub>54</sub>O<sub>6</sub> |
| Formula weight | 630.86 |
| Temperature/K | 296 (2) |
| Crystal system | monoclinic |
| Space group | C2 |
| a/Å | 19.1677(11) |
| b/Å | 11.5860(7) |
| c/Å | 16.8399(11) |
| α/° | 90 |
| β/° | 90 |
| γ/° | 90 |
| Volume/Å<sup>3</sup> | 3739.8(4) |
| Z | 110 |
| ρ<sub>calc</sub>/g/cm<sup>3</sup> | 1.417 |
| μ/mm<sup>-1</sup> | 0.131 |
| F(000) | 1650 |
| Crystal size/mm<sup>3</sup> | 0.35 × 0.18 × 0.15 |
| Radiation | MoKα (λ = 0.71073) |
| 2θ range for data collection/° | 0.995 to 25.75 |
| Index ranges | -23 ≤ h ≤ 23, -14 ≤ k ≤ 14, -20 ≤ l ≤ 20 |
| Reflections collected | 108274 |
| Independent reflections | 108274 [R<sub>int</sub> = 0.0811, R<sub>sigma</sub> = 0.0325] |
| Data/restraints/parameters | 7123/3/415 |
| Goodness-of-fit on F<sup>2</sup> | 1.045 |
| Final R indexes [I ≥ 2σ (I)] | R<sub>1</sub> = 0.1038, wR<sub>2</sub> = 0.2797 |
| Final R indexes [all data] | R<sub>1</sub> = 0.1362, wR<sub>2</sub> = 0.3157 |

### Chromatin Immunoprecipitation

Nuclear ChIP was performed using intact nuclei from cells after removal of lipids. 3T3L1 that were either undifferentiated or differentiated for 3 to 6 days were cross-linked using 1% formaldehyde. Nuclear fractions were obtained by sequentially lysing the cells in two buffers-lysis buffer 1 (140 mM NaCl, 1 mM EDTA, 50 mM HEPES, 10% Glycerol, 0.5% NP-40, 0.25% Triton X-100) and lysis buffer 2 (10 mM Tris pH 8, 200 mM NaCl, 1 mM EDTA, 0.5 mM EGTA). This was followed by lysis in SDS lysis buffer (1% SDS, 10 mM EDTA, 50 mM Tris-HCl pH 8). Lysates were sonicated in a Diagenode Bioruptor (Liège, Belgium) to produce 200 to 1000 bp DNA fragments. The cell lysates were incubated with H4K5 Bu antibody (PTM Biolabs) or preimmune IgG per sample and 25 μl of magnetic protein G Dynabeads (catalog no. 10003D; Novex, Waltham, MA) overnight at 4°C. Beads were then washed successively with low-salt buffer (0.1% SDS, 1% Triton X-100, 2 mM EDTA, 20 mM Tris-HCl pH 8, and 150 mM NaCl), high-salt buffer (0.1% SDS, 1% Triton X-100, 2 mM EDTA, 20 mM Tris-HCl pH 8, and 500 mM NaCl), LiCl buffer (250 mM LiCl, 1% NP-40, 1% NaDOC, 1 mM EDTA, and 10 mM Tris-HCl, pH 8), and TE (10 mM Tris-HCl pH 8 and 1 mM EDTA). DNA-protein complexes were recovered from beads in elution buffer (0.1% SDS and 100 mM NaHCO3). The ChIP eluates and input samples in the elution buffer were then reverse cross-linked by adding 200 mM NaCl and 20 μg proteinase K (Sigma) and incubating at 65°C for 4 h. Subsequently, 20 μg of RNase A (Sigma) were added and the samples were further incubated for 15 min at 37°C. The immunoprecipitated DNA was extracted using phenol-chloroform, ethanol precipitated, and used for quantitative PCR. The region-specific primer sets used for the ChIP-qPCR analysis have been mentioned in Table 3. These primers were designed based on the genomic coordinates with enriched H3K27 acetylation in the vicinity of promoters of candidate genes in brown adipose tissue, the information for which was obtained from NCBI37/mm9 assembly (ENCODE/LICR).

### Immunohistochemistry

Collected liver tissue samples were stored in 4% paraformaldehyde for 24 hours after which they were cryoprotected in 30% sucrose solution for two weeks. Cryosections were performed at 7 µm sections using Cryostat LeicaCM1850 UV. For staining, the tissue sections were washed with PBS followed by antigen retrieval with 0.01 M citrate buffer (pH 6). The tissues were then permeabilised with 0.3% Triton X-100/PBS (PBST) and blocked with 2% serum followed by incubation with primary antibodies overnight at 4ᴼC. The next day, secondary antibody incubation was carried out for one hour at room temperature followed by staining of the nuclei with Hoechst 33342 and mounting with 70% glycerol. Images were acquired by confocal microscopy. For quantitation, the intensity of each modification-specific staining was normalized with respect to the Hoechst staining intensity.

### Reagents and antibodies

The media used for cell culture was purchased from Sigma-Aldrich; DMEM (Ref: 1152). Several other cell culture reagents were obtained from HIMEDIA, such as PBS (Ref: TL1006), Trypsin-EDTA solution 10X (Ref: TCL070) and Antibiotic antimyotic solution 100X (Ref: A002A). Multiple reagents used for cell culture were obtained from Sigma-Aldrich: Insulin (Ref: I6634), Dexamethasone (Ref: D4902), IBMX (Ref: I5879). TRIzol reagent was obtained from Ambion life technologies (Ref: 15596018). The commercial antibodies used in this study are Pan anti-butyryllysine (PTM Biolabs, SKU: PTM 301), Butyryl-Histone H4 (Lys 12) rabbit pAb (PTM Biolabs, SKU: PTM 308), Butyryl-Histone H3 (Lys 9) rabbit pAb (PTM Biolabs, SKU: PTM 305), Butyryl-Histone H3 (Lys 23) mouse mAb (PTM Biolabs, SKU: PTM 307), Butyryl-Histone H4 (Lys 8) rabbit pAb (PTM Biolabs, SKU: PTM 311), Butyryl-Histone H4 (Lys 5) rabbit pAb (PTM Biolabs, SKU: PTM 313), Butyryl-Histone H3 (Lys 27) rabbit pAb (PTM Biolabs, SKU: PTM 315).

### Statistical analysis

All statistical analyses were performed using GraphPad Prism 7 Software (California, USA). Data obtained from two or three individual experiments as mentioned in figure legends, were expressed as mean +/- SD. Either two-tailed unpaired Student’s t-test or one-way ANOVA with Dunnett’s/Bonferroni’s multiple comparision was used to determine the statistical significance values.

The mutagenecity analysis data were analyzed by comparing the groups using one-way ANOVA followed by Neman-Keuls test. Body weight data for acute toxicity test were summarized in Mean ± SD and compared by two factor (Group and Period) repeated measure analysis of variance (RM ANOVA) and the significance of mean difference within (intra) and between (inter) the groups was done by Newman-Keuls post hoc test. A P-value of equal to or less than 0.05 was considered statistically significant.

## Acknowledgements

TKK is a recipient of JC Bose fellowship from the Department of Science and Technology, Govt. of India (SR/S2/JCB-28/2010). AB is a senior research fellow of UGC. This work was supported by Life Science Education & Training at JNCASR grant (DBT/INF/22/SP27679/2018). We acknowledge Dr. Rahul Gajbhiye from NIPER, Kolkata for the assistance in mass spectrometry-related experiments in their central instrument facility. We thank Dr. R.G. Prakash for providing the necessary facilities to perform the mice experiment in Animal Facility, JNCASR and the confocal facility, JNCASR for the immunofluorescence imaging analysis. We also thank Kruthi HT for helping in the preparation of the figures and graphical abstract. We further acknowledge the Sophisticated Analytical Instrument Facility and Research (SAIFR, Lucknow) for providing spectral data. This manuscript bears CDRI communication number 2021/TK.

## Author Contributions

AB and TKK conceived the project. AB designed and performed all the biological experiments. SC performed the initial synthesis of all compounds and molecular docking analysis. KVS performed bulk scale synthesis of LTK-14A for animal experiments. AKS and RG assisted in the animal experiments. UB and MV performed *in silico* analysis of high throughput RNA sequencing data. AN did the toxicity evaluation and SKR evaluated the mutagenecity. NN provided the raw material *Garcinia indica* for compound isolation. AB and TKK analyzed the data and wrote the manuscript.

## Results

### Physiological levels of butyryl CoA and histone butyrylation increase during adipogenesis

Under normal circumstances, the cellular levels of specific coenzyme A derivatives are much less than acetyl CoA (6). We surmised that the situation might alter in particular cases where metabolic pathways promoting the production of other acyl CoA derivatives become more predominant. One such possible scenario could be the process of adipogenesis in which fatty acid synthesis and breakdown can give rise to several intermediate products like butyryl CoA.

The causal relationship between butyryl CoA and adipogenesis was investigated using the 3T3L1 pre-adipocyte cell line as a model system (fig 1A). Our RT-qPCR data showed a significant increase in the expression of *Acss2* (acyl-CoA synthetase short-chain family member 2) upon differentiation (fig 1B). Previously, ACSS2 had been established as the enzyme responsible for the generation of acyl-CoA derivatives from their corresponding short chain fatty acids (7). Indeed, siRNA mediated knockdown of *Acss2* (fig 1C) led to a drastic reduction in histone butyrylation with a relatively lesser effect on acetylation (fig 1D). This indicated that the ACSS2 generated butyryl CoA might be playing an important physiological role in adipogenesis. These findings prompted us to investigate whether adipocyte differentiation also led to an increased butyrylation of proteins. We observed that in accordance with the previous report (4), there was an increase in acetylation level at histone H3 lysine 9 (H3K9 Ac) in the differentiated cells as compared to the undifferentiated cells (Fig. 1C, panel i). Interestingly immunoblotting with pan butyryllysine and site-specific antibodies (H3K9, H3K23, H3K27, H4K5, H4K8 and H4K12) against butyrylation (Fig 1C, panel ii and iii) revealed that butyrylation at each of these sites also increased upon differentiation. Chromatin immunoprecipitation with antibody against H4K5 butyrylation revealed that the modification enrichment levels increased in promoters of different pro-adipogenic genes in a time dependant manner. *Pparg*, the master regulator of adipogenesis, exhibited a steady increase in butyrylation level in its promoter with the progress of differentiation (fig 1F, panel i) while butyrylation increased only at the terminal stage of adipogenesis in the promoter of *Lep*, encoding the adipocyte marker leptin which gets expressed in mature adipocytes (fig 1F, panel ii). Interestingly, the promoter of *Cebpd*, which regulates *Pparg* expression did not show any enhanced butyrylation in its promoter on day 3 and instead, exhibited increased butyrylation on day 6 of adipogenesis (fig 1F, panel iii). Collectively these results showed that during adipogenesis, EP300 catalysed butyrylation of histones could be an important epigenetic signature.

**Fig. 1.**
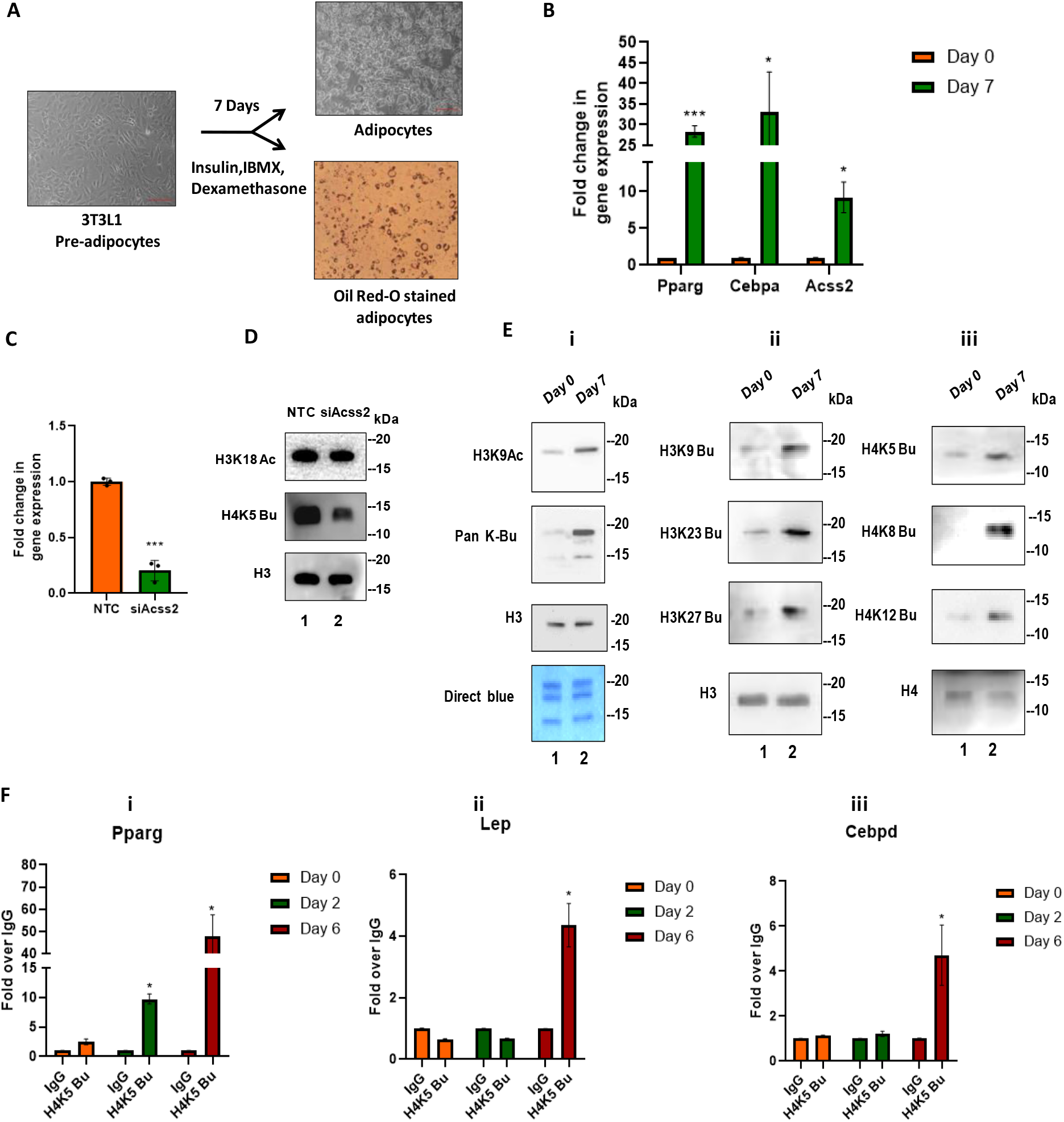
Histone butyrylation increases during adipogenesis. (A) Phase contrast images of 3T3L1 cells exhibiting fibroblast like morphology in their pre-adipocyte state and accumulation of lipid droplets in their adipocyte state. The lipid droplets can be stained using Oil Red-O dye. (B) RT-qPCR analysis was performed to check for transcript levels of Acss2 in undifferentiated and differentiated 3T3L1 cells. Error bars denote mean +/-SEM of three biological replicates; two-tailed unpaired Student’s t-test: * P < 0.05, **P < 0.01, ***P < 0.001, ns: not significant. (C) Knockdown of Acss2 in differentiated 3T3L1 cells was confirmed by RT-qPCR analysis. Error bars denote mean +/- SEM of three biological replicates; two-tailed unpaired Student’s t-test: * P < 0.05, **P < 0.01, ***P < 0.001, ns: not significant. (D) Histone acetylation and butyrylation pattern in 3T3L1 cells upon Acss2 knockdown was checked using antibodies against acetylated histone H3K18 and butyrylated H4K5. (E) Histone acetylation and butyrylation levels in 3T3L1 cells in undifferentiated and differentiated state were analysed by immunoblotting with antibodies specific for acetylated H3K9, butyrylated H3K9, H3K23, H3K27, H4K5, H4K8, H4K12 and pan-butyryl lysine antibodies. Antibodies against core histones H3 and H4 were used as loading controls along with Direct Blue staining of all four core histones. Lane 1-undifferentiated pre-adipocyte (Day 0) and Lane 2 -differentiated adipocyte (Day 7) in each of the panels i, ii and iii. (F) Enrichment of H4K5 butyrylation in the promoters of (i) Pparg, (ii) Lep and (iii) Cebpd represented in the form of fold change over IgG. Error bars denote mean +/- SEM of three biological replicates; one-way ANOVA with Dunnett’s multiple comparision was performed: * P < 0.05, **P < 0.01, ***P < 0.001, ns: not significant.

### Identification of a small molecule that selectively inhibits p300 catalysed histone butyrylation *in vitro*

In order to investigate the contribution of p300 catalyzed butyrylation per se in adipogenesis, we adopted a chemical biology-based approach. For this purpose, a series of derivatives of garcinol, a polyisoprenylated benzophenone isolated from *Garcinia indica*, were synthesized (8). These derivatives, unlike garcinol, were found to be non-toxic to cells and showed a high specificity toward the catalytic activity inhibition of EP300. The derivatives were generated by internal cyclisation of garcinol with further addition of one or two functional groups to the remaining free -OH groups-giving rise to mono and disubstituted derivatives (Sup. fig. 1A). Amongst these, a few monosubstituted derivatives could potently inhibit EP300 catalysed acetylation as well as butyrylation, while their disubstituted counterparts were unable to inhibit acetylation (Sup. fig. 2 B, C and D). This was probably due to increased steric hindrance brought about by the addition of an extra functional group that interfered with the interaction between the compounds and EP300.

An inverse correlation between catalytic rates of acylation reactions by EP300 and increasing acyl chain length had already been established previously (9). Based on this, we hypothesized since butyrylation is catalysed by EP300 at a much slower rate than acetylation, one of the compounds that could not inhibit EP300 catalysed acetylation might be able to inhibit the slower butyrylation reaction. Our *in vitro* butyrylation assay using recombinant p300 and histone H3 as the substrate revealed a few potential candidates (Sup. Fig. 2 E) as butyrylation inhibitors. Amongst them, LTK-14A, which is 13, 14 dimethoxy isogarcinol, was capable of inhibiting full-length EP300 catalysed butyrylation at a concentration of 25 μM (fig. 2 C and D), with minimal effect on EP300 catalysed acetylation in similar reaction conditions (fig. 2 A and B).

**Figure 2:**
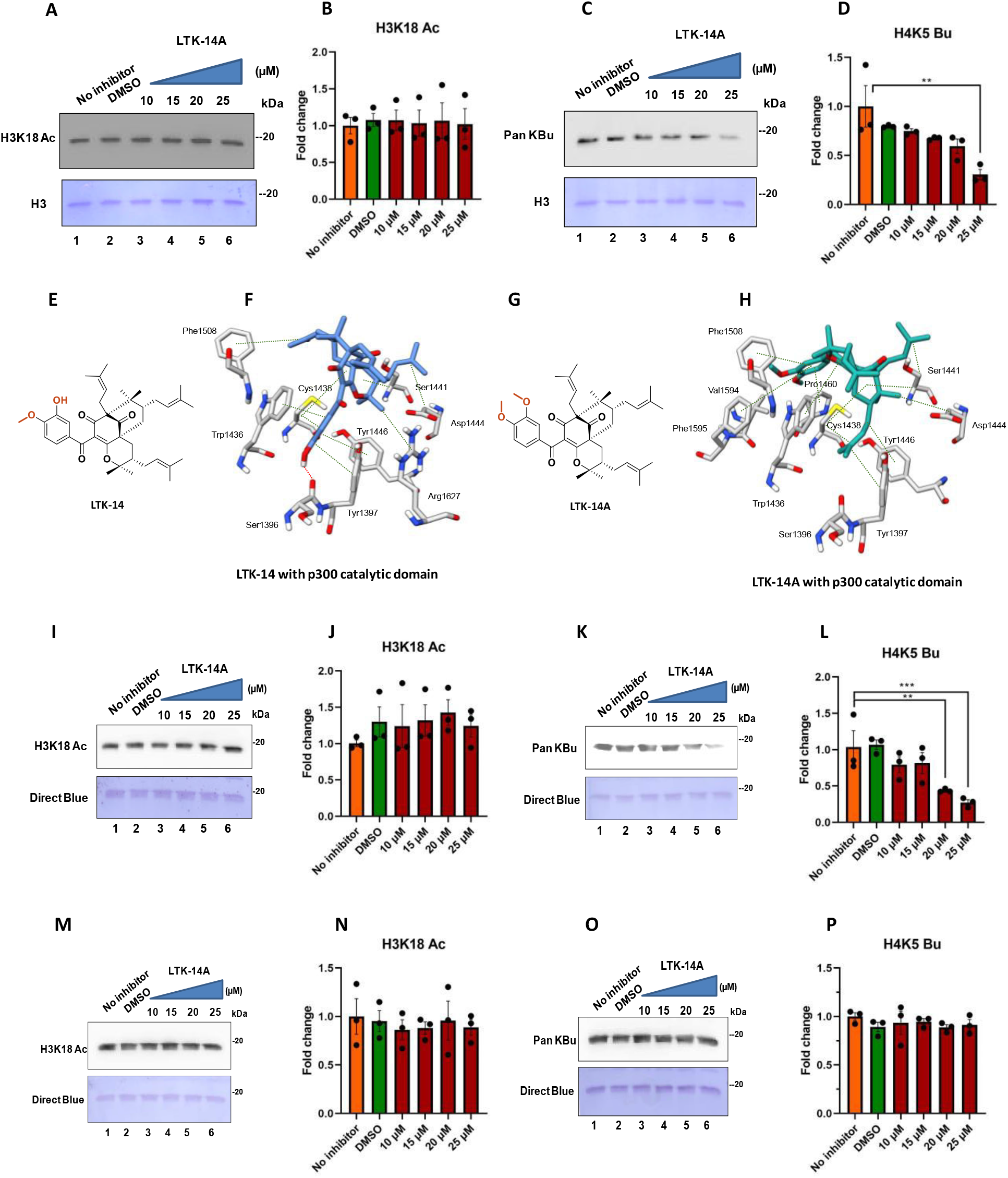
LTK-14A selectively inhibits p300 catalysed butyrylation without affecting its acetyltransferase activity: Immunoblotting with antibodies against acetylated H3K18 (A) and pan-butyryl lysine (C) to check for the relative inhibition of acetyltransferase and butyryltransferase activities of full length p300 by LTK-14A at different concentrations. Recombinant histone H3 was used as substrate. Lane 1, enzyme alone; Lane 2, enzyme + DMSO; Lane 3-6, enzyme + LTK-14A at concentrations 10, 15, 20 and 25 µM respectively. Images of Western blotting are given in panels F and H and their quantitations are given in panels B and D. Molecular structures of mono-(LTK-14) and di-(LTK-14A) substituted isogarcinol (E and G). Molecular docking studies for studying comparative interaction of LTK-14 and LTK-14A with the amino acid residues of p300 catalytic domain (F and H). The predicted interactions are denoted by tick lines. Hydrogen bonding is shown in red and hydrophobic interaction is shown in green. Immunoblotting with antibodies against acetylated H3K18 (I) and pan-butyryl lysine (K) to check for the relative inhibition of acetyltransferase and butyryltransferase activities of wild type p300 catalytic domain by LTK-14A at different concentrations. Lane 1, enzyme alone; Lane 2, enzyme + DMSO; Lane 3-6, enzyme + LTK-14A at concentrations 10, 15, 20 and 25 µM respectively. Images of Western blotting are given in panels I and K and their quantitations are given in panels J and L. Immunoblotting with antibodies against acetylated H3K18 (M) and pan-butyryl lysine (O) to check for the relative inhibition of acetyltransferase and butyryltransferase activities of catalytically active p300 catalytic domain with point mutation C1438A by LTK-14A at different concentrations. Lane 1, enzyme alone; Lane 2, enzyme + DMSO; Lane 3-6, enzyme + LTK-14A at concentrations 10, 15, 20 and 25 µM respectively. Images of Western blotting are given in panels M and O and their quantitation are given in panels N and P. Error bars denote mean +/- SEM of three biological replicates, one-way ANOVA with Bonferroni’s multiple comparison; * P < 0.05, **P < 0.01, ***P < 0.001, ns: not significant.

In order to understand the underlying mechanism of the selective inhibition by LTK-14A, we performed molecular docking analysis of LTK-14A (Fig. 2 G) and the catalytic domain. To delineate this, we solved the crystal structure of LTK-14A (Table 1, Sup. Fig.2A). LTK-14 (fig. 2E), a specific inhibitor of EP300 catalysed acetylation and butyrylation (Sup. Fig. 2 B and C) was used for further comparative analysis. The energy minimized analysis revealed that both LTK-14 and LTK-14A could potentially bind with the active site of the EP300 catalytic domain forming several hydrophobic interactions (Fig.2 F and H). LTK-14, due to the presence of a free phenolic –OH group, could hydrogen bond with a carbonyl oxygen atom present in the peptide backbone of S1396 residue. However, LTK-14A has all of its phenolic –OH groups blocked and so it is incapable of forming such an interaction. This hydrogen bonding by LTK-14 could imply a stronger binding to the enzyme, while LTK-14A, by interacting with fewer residues presumably inhibits only the slower butyrylation reaction and the interaction may not be strong enough to deter the faster acetylation reaction.

Some of the residues with which LTK-14A could potentially form hydrophobic interaction have already been reported before as being critical for targeting inhibition of the enzymatic activity of EP300 by the first reported EP300 inhibitor lysyl CoA (10). These residues include C1438, Y1446, Y1397 and W1436. To verify this observation we generated a point mutant of p300 catalytic domain in which cysteine 1438, showing the highest propensity to interact with LTK-14A, was mutated to alanine. We found that the C1438A catalytic domain is equally active as the wild-type catalytic domain in an *in vitro* lysine acetyltransferase assay (Sup. Fig. 2B). We observed that the wild-type EP300 catalytic domain catalysed butyrylation was inhibited by LTK-14A most prominently at 25 µM concentration while acetylation was not inhibited (compare lane 6 in each of fig.2 I-L). In contrast, neither acetylation nor butyrylation modification catalysed by the mutant EP300 catalytic domain could be inhibited by LTK-14A even at 25 µM concentration (compare lane 6 in each of fig. 2 M-P). Altogether, these data suggest that LTK-14A indeed interacts with C1438 within the active site of the EP300 catalytic domain, and this residue is critical for its interaction to bring about butyrylation reaction inhibition.

### LTK-14A is a metabolically stable, cell-permeable compound that significantly inhibits adipogenesis

Upon establishing that LTK-14A is an EP300 acylation-specific inhibitor *in vitro*, its cellular effects were tested using 3T3L1 cells. The quality of our compound used in all biological experiments was acceptably good with more than 97% purity (Sup. Fig. 2 C, D and E). We found the LTK14A is highly cell-permeable and metabolically stable as it could be detected by mass spectrometry analysis from metabolites extracted from LTK-14A treated cells after six days (Fig. 3A). MTT assay showed that LTK-14A at 25 μM concentration did not have any adverse effect on the cellular oxidoreductase activity, a signature of metabolic output of cells, (Fig. 3B) suggesting that LTK-14A is non-toxic to 3T3L1 even after 6 days of treatment. Oil Red-O staining of 3T3L1 cells differentiated in the presence of LTK-14A followed by spectrophotometric quantitaion revealed significant inhibition of adipogenesis by the compound (Fig. 3C, D).

**Fig. 3:**
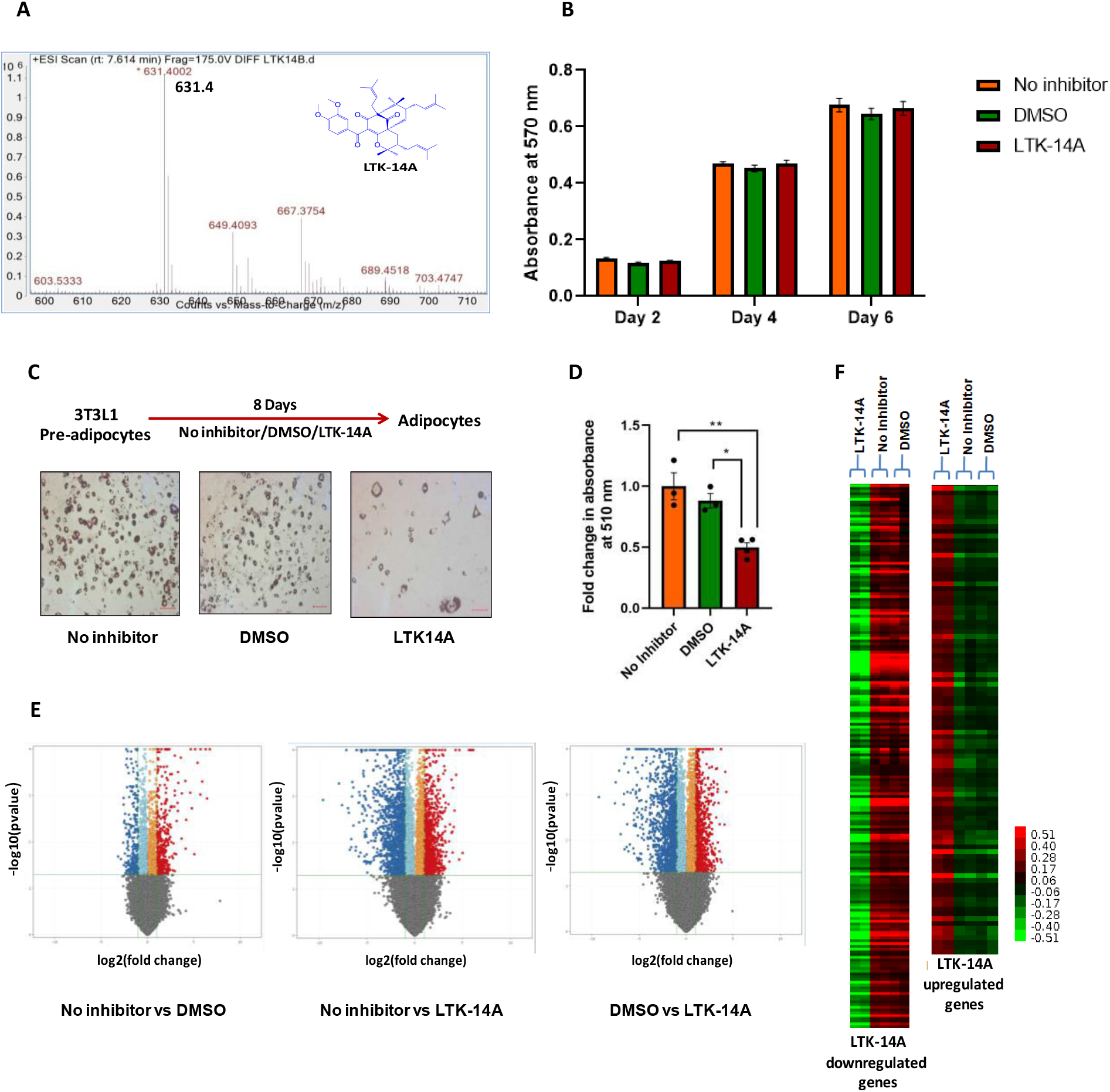
The butyrylation inhibitor represses adipogenesis in 3T3L1 cells: (A) The permeability and stability of LTK-14A in 3T3L1 was tested by LC-ESI-MS analysis of metabolites, extracted from the compound treated 3T3L1 cells to detect the protonated LTK-14A in the ESI spectra of the eluted fraction. (B) MTT assay was performed to check for any possible cellular toxicity effect of LTK-14A, using 25 µM concentration of the compound on 3T3L1 cells upon incubation for varying lengths of time (2, 4 and 6 days). Comparison was done with cells that were left untreated or treated with DMSO as solvent control. The effect of LTK-1a4A (25 µM) treatment on adipogenesis was tested by Oil Red-O staining of the adipocytes (C) and the quantitation of staining is represented graphically in panel D. Error bars denote mean +/- SEM of three biological replicates, one-way ANOVA with Bonferroni’s multiple comparison; * P < 0.05, **P < 0.01, ***P < 0.001, ns: not significant. (E) The effect of LTK-14A on the global transcription network inside 3T3L1 cells was investigated by high throughput transcriptome analysis with RNA isolated from compound treated, DMSO treated and untreated 3T3L1 cells (n=2 in each case). Differential gene expression pattern of three conditions compared with each other, depicted in volcano plots-No Inhibitor versus DMSO, No Inhibitor versus LTK-14A and DMSO versus LTK-14A. The downregulated and upregulated genes are shown in blue and red respectively.(F) Heat maps showing the differential gene expression pattern between LTK-14A treated cells and DMSO treated or untreated cells.

In order to understand the transcriptional mechanism of LTK14A-mediated butyrylation inhibition on adipogenesis, adipogenic cells were treated with LTK-14A and subjected to transcriptome analysis. We used untreated and solvent treated cells as control. Volcano plot (Fig. 3E) and heat map analysis (Fig. 3F) revealed that upon treatment with the compound, a large number of genes were differentially expressed compared to DMSO treated and untreated conditions. To better understand the LTK-14A affected genes, we focused on the common genes that were differentially expressed in the two cases-(1) LTK-14A versus no inhibitor and (2) LTK-14A versus DMSO. Among these common genes, 2200 genes were found to be downregulated and 1309 genes were upregulated (Fig. 4A) with a majority of them to be protein-coding genes (Fig. 4B). Gene ontology analysis revealed that LTK14A treatment predominantly affected the genes involved in lipid metabolism pathways (Fig. 4C, E) which were minimally affected by DMSO alone (Fig. 4D). The gene regulatory networks and the respective pathways were constructed with the major genes that were affected by LTK-14A treatment compared to DMSO treated and untreated conditions (Fig. 4F, Sup. Fig.3A). They collectively depict how LTK-14A, by inhibiting the expression of a large number of genes that are interconnected via several pathways associated with lipid metabolism, brings about a collapse in the adipogenesis process as a whole.

**Figure 4:**
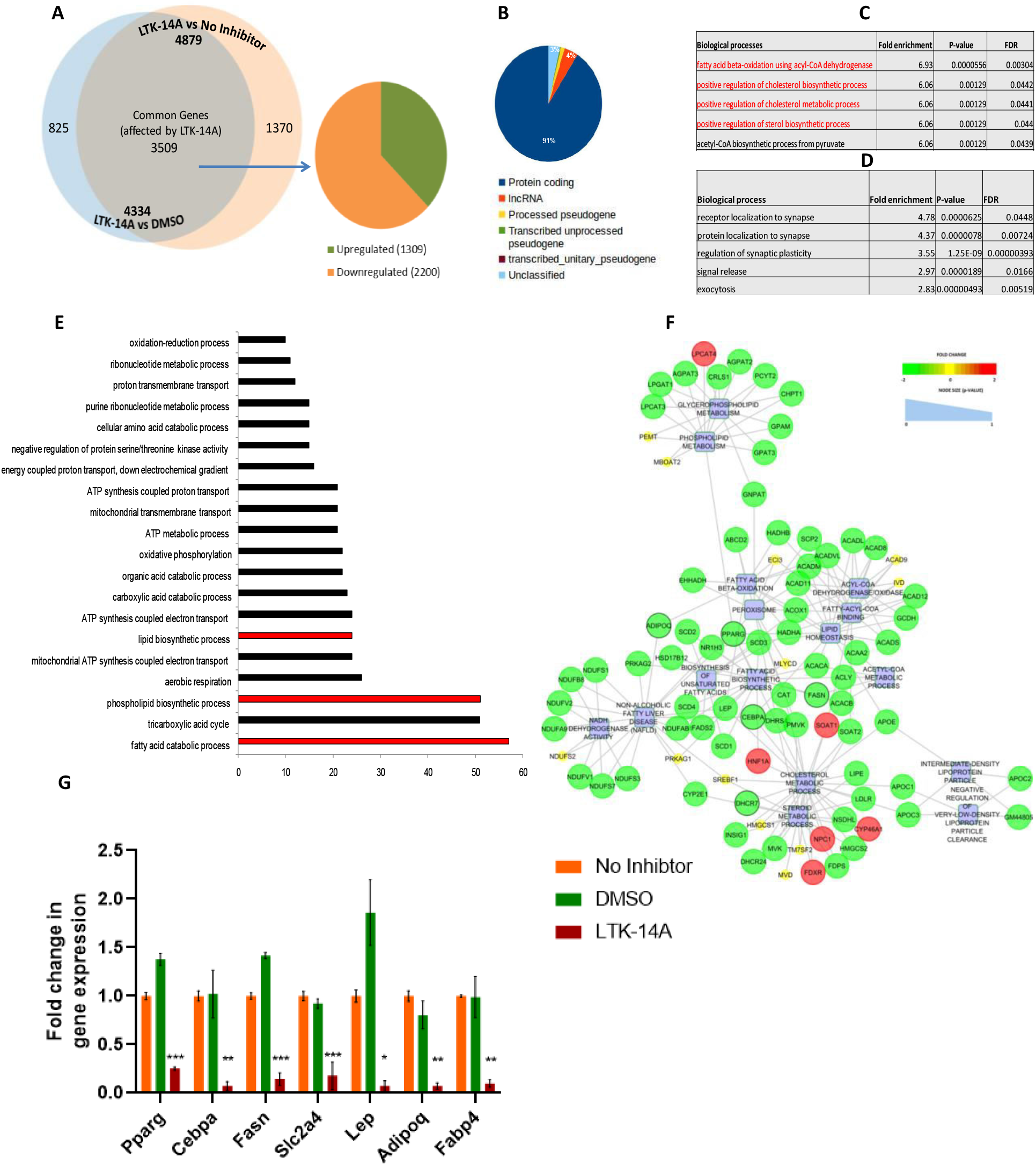
LTK-14A represses expression of several pro-adipogenic genes and inhibits lipid metabolism: (A) Venn diagram showing the number of differentially expressed genes upon LTK-14A treatment with respect to untreated and DMSO treated conditions. (B) Classification of the common differentially expressed genes according to their type. Gene ontology analysis was performed to study the different biological pathways that were affected by LTK-14A in 3T3L1 compared to DMSO treated as well as untreated conditions, as predicted by Panther software. The pathways were ranked in the order of those with maximum number of genes affected (E). A different ranking order was created on the basis of fold enrichment i.e. the pathways in which the ratio of number of observed affected genes over number of expected genes to be affected was the highest (C). A similar analysis was also performed for genes differentially expressed in DMSO versus untreated conditions (D). (F) A molecular network was constructed to depict the interconnectivity between the lipid metabolism related genes and pathways that were affected upon their dysregulation by LTK-14A compared to DMSO treated conditions. (G) RT-qPCR analysis was performed to validate the putative target genes of LTK-14A. Error bars denote mean +/- SEM of three biological replicates; one-way ANOVA with Bonferroni’s multiple comparison: * P < 0.05, **P < 0.01, ***P < 0.001, ns: not significant.

A few of the pro-adipogenic genes downregulated by LTK-14A, as indicated by RNA sequencing, were selected for further validation (Sup. Fig. 3B). RT-qPCR analysis corroborated that except for *Cebpd*, most of the other pro-adipogenic genes (*Pparg, Cebpa, Adipoq, Lep, Slc2a4, Fasn* and *Fabp4*) showed a marked downregulation upon LTK-14A treatment (Fig. 4G). *Acss2* was also downregulated by LTK-14A (Sup. Fig.4C). The upregulation of anti-adipogenic genes was also verified by performing RT-qPCR analysis for two such candidate genes- *Kdm4c* and *Tead4* (Sup. Fig.3C). Collectively, the RNA sequencing analysis and the validation of gene targets further convincingly suggested that LTK-14A can inhibit adipogenesis in 3T3L1 cells, mainly by repressing the expression of important pro-adipogenic genes.

### LTK-14A inhibits histone butyrylation in 3T3L1 cells, without affecting acetylation or other post-translational modifications

Furthermore we wanted to find out whether LTK-14A mediated downregulation of adipogenesis is causally linked to the inhibition of histone butyrylation in the 3T3L1 cells. We observed that treatment with LTK-14A inhibited overall butyrylation of histones (Fig. 5 A, B). Among the specific modification sites tested, H3K23 and H4K5 butyrylation showed a significant decrease upon treatment with LTK-14A (Fig. 5 A, B) while butyrylation at other tested sites was affected minimally (Sup. Fig. 3D). Acetylation of histones (H3K9, H3K18 and H4K12) was also found to be unaffected (Fig. 5 C, D) upon LTK-14A treatment. To establish the specificity of the LTK14A, the effect of LTK-14A on lysine and arginine methylation along with phosphorylation of histones was also investigated. We observed (Fig. 5 E, F) no significant changes in these marks upon the compound treatment, thereby indicating that in the 3T3L1 cells, LTK-14A specifically inhibits butyrylation.

**Fig. 5:**
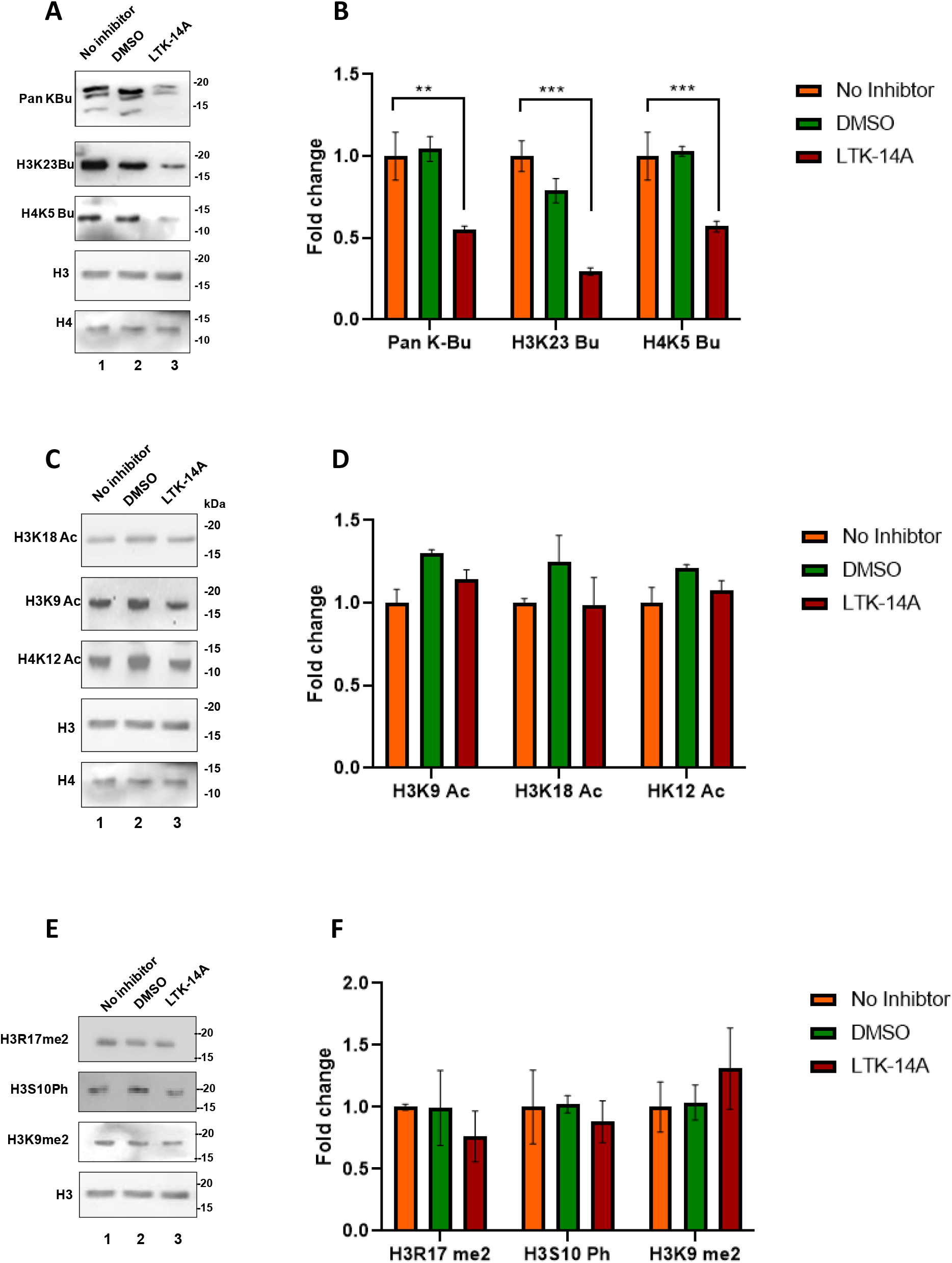
LTK-14A inhibits histone butyrylation in 3T3L1 cells, without affecting acetylation or other post-translational modifications: Histone butyrylation levels in 3T3L1 cells that were treated with LTK-14A were compared with that in DMSO treated and untreated conditions by immunoblotting with antibodies against butyrylated histone H3K23, H4K5 and pan-butyryl lysine. Histones H3 and H4 were used as loading controls (A). The fold change of quantified band intensity is graphically represented in panel (B) in which the error bars denote mean +/- SEM of three biological replicates. Under similar conditions histone acetylation levels were compared by immunoblotting using antibodies against acetylated histones H3K18, H3K9 and H4K12 (C). The result is graphically represented in panel (D) in which error bars denote mean +/- SEM of three biological replicates. Immunoblotting was also performed with antibodies against histone H3R17 dimethylation, H3S10 Phosphorylation and H3K9 dimethylation (E). The result is graphically represented in panel F in which error bars denote mean +/- SEM of three biological replicates. For all data one-way ANOVA with Bonferroni’s multiple comparision was performed: * P < 0.05, **P < 0.01, ***P < 0.001, ns: not significant.

### Inhibition of butyrylation attenuates weight gain of high fat diet fed C57BL6/J mice

After observing the anti-adipogenic effect of LTK-14A on 3T3L1 cells, we investigated whether this compound could also affect adipogenesis and obesity at an organismal level. For this purpose, C57BL6/J mice were maintained either on a normal diet or a high fat diet with or without LTK-14A. In due course of time it was observed that the mice maintained on a high fat diet along with LTK-14A had a weight gaining trend intermediate between the normal diet and high fat diet fed mice (Fig.6A,B). Dissection of the mice revealed that the fat accumulation in high fat diet-fed mice was the highest, followed by those on high fat diet and LTK-14A, and then by those on normal diet (Fig. 6C). The epididymal fat pads were separately weighed and again it was observed that the weights of the fat pads from compound treated mice were much less than that of high fat diet-fed mice (Fig. 6 D, E). The diet consumption pattern of the mice showed that the two groups on high fat diet with or without LTK-14A, had a similar consumption rate indicating that the reduced weight gain of the compound treated mice is not due to discrepancy in diet consumption (Table 2). Hematoxylin and eosin staining of adipose tissue and liver (Fig. 6 F,G) showed signs of adipocyte hypertrophy and hepatic steatosis in high fat diet-fed mice, which were absent in the mice maintained on normal diet or high fat diet with LTK-14A.

**Fig. 6:**
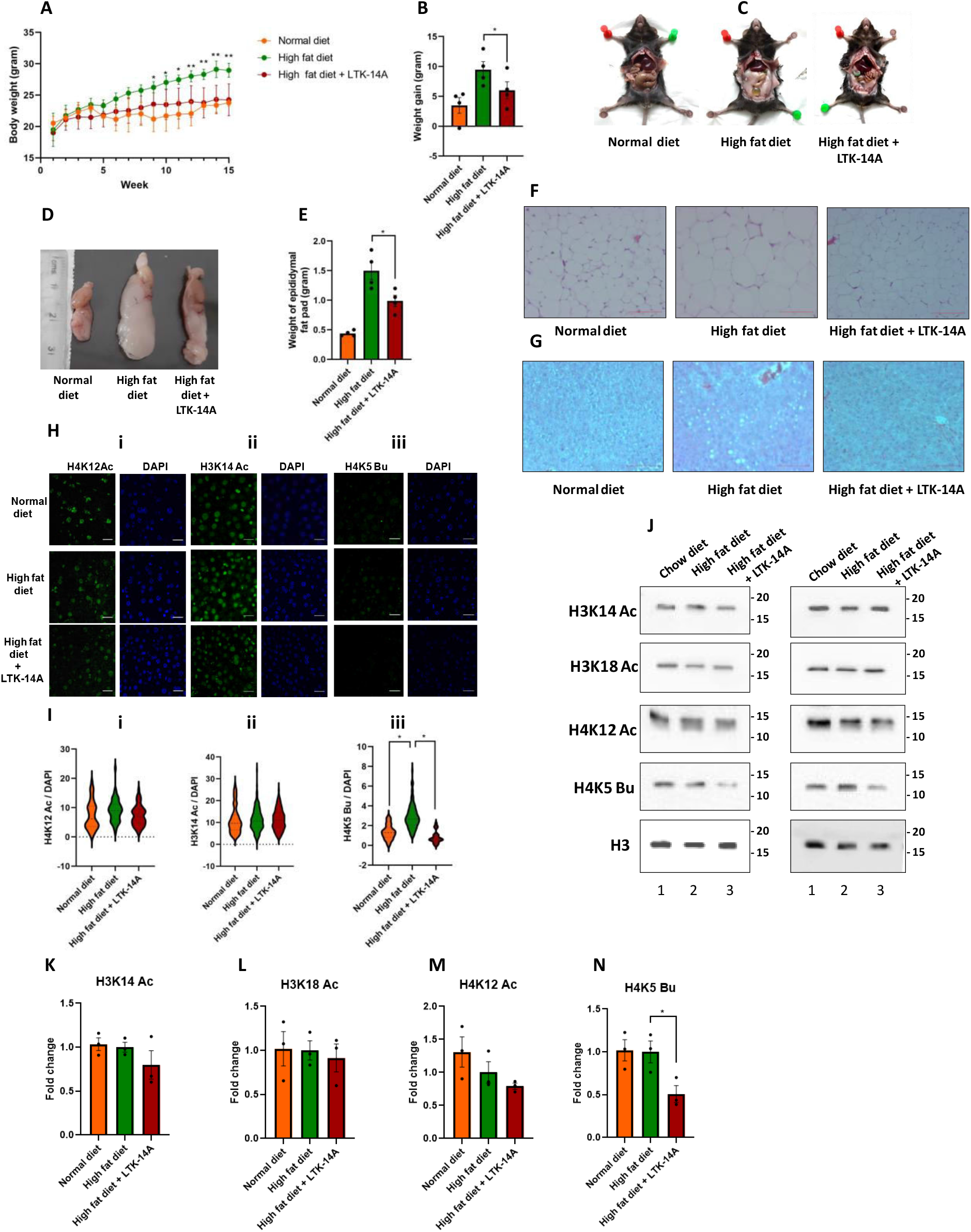
The weight gaining of high fat diet fed mice (C57BL6/J) could be significantly reduced by the butyrylation specific inhibitor of p300: Trend of weight gaining (A) and average body weight gained after 16 weeks (B) of C57BL6/J mice maintained on normal chow diet, high fat diet and high fat diet mixed with LTK-14A was plotted; n=4 in each group. Representative images of dissected bodies showing fat accumulation (C) and epididymal fat pads (D) of the mice from three groups. The average weight of the fat pads is represented graphically (E); n=4 in each group. Representative images of morphology of adipose tissue (F) and liver (G) from the mice of the three groups, as studied by hematoxylin and eosin staining. For all data one-way ANOVA with Dunnett’s multiple comparision was performed: * P < 0.05, **P < 0.01, ***P < 0.001, ns: not significant. (H) Immunofluorescence microscopy of liver sections stained with antibodies against histone H4K12 acetylation (panel i), histone H3K14 acetylation (panel ii) and histone H4K5 butyrylation (panel iii). The fluorescence intensity of the modified histone marks was normalised against DAPI staining of whole nuclei for quantitation which is graphically depicted in (I) panel i, ii and iii ;n=4 in each group; one-way ANOVA with Dunnett’s multiple comparision: * P < 0.05, **P < 0.01, ***P < 0.001, ns: not significant. (J) Histone H3K14, H3K18, H4K12 acetylation and H4K5 butyrylation levels in epididymal fat pads of mice were estimated by immunoblotting. Histone H3 was used as loading control The fold change of quantified band intensity is graphically represented in panel K, L, M and N ;n=3 in each group; one-way ANOVA with Dunnett’s multiple comparision: * P < 0.05, **P < 0.01, ***P < 0.001, ns: not significant.

Immunofluorescence staining of the liver tissue samples revealed that there was a significant increase in H4K5 butyrylation levels in high fat diet-fed mice compared to normal diet-fed mice in which the modification was barely detectable and it was almost completely inhibited upon LTK-14A treatment (Fig. 6H panel iii, Fig. 6I panel iii). On the other hand, no significant difference in H4K12 (Fig. 6H panel i, Fig. 6I panel i) and H3K14 acetylation (Fig. 6H panel ii, Fig. 6I panel ii) levels could be observed across all the three groups. Furthermore, histone modification pattern in the adipose tissues was studied with acid extracted histones from the epididymal fat pads. H4K5 butyrylation was reduced in the LTK-14A treated samples compared to those of high fat diet-fed mice (Fig. 6 J, N) while H3K14, H3K18 and H4K12 acetylation levels were relatively unchanged across all the three groups (Fig. 6 J, K, L, M). The metabolic stability of LTK-14A in mice was verified by its detection in ESI spectra of analytes extracted from the liver of compound treated mice (Sup. Fig 3 E).

**Table 2:** Food consumption pattern of mice used in obesity studies

|  |  |  |  |
| --- | --- | --- | --- |
| **Set A** |  |  |  |
| **Type of diet** | **Chow diet** | **High fat diet** | **High fat diet and LTK-14A** |
| Average food consumed per mice per day (gram) | 8.0157 | 1.95547 | 2.04051 |
| Standard error of mean | 1.31838 | 0.23142 | 0.24589 |
| **Set B** |  |  |  |
| **Type of mice** | **db +/-** | **db/db and vehicle control** | **db/db and LTK-14A** |
| Average food consumed per week per mouse (gram) | 25.8 | 37.8 | 34 |
| Standard error of mean | 1.04 | 1.64 | 2.08 |

### The butyrylation specific inhibitor reduces the weight of genetically obese db/db mice

The previous model system involved the use of LTK-14A as a prophylactic. To test whether LTK-14A could also reduce obesity by inhibiting histone butyrylation, genetically obese homozygous mutant db/db mice were used. Heterozygous mutant db+/- mice were taken as control. The hyperphagic db/db mice were orally administered LTK-14A daily at the same dosage as before (50 mg/Kg body weight). We observed that upon oral gavaging of LTK-14A, the overweight mice not only showed an arrest in weight gain, but their body weights were significantly reduced by the end of one month of treatment (Fig. 7 A, B, C). The food intake of LTK-14A treated mice was marginally less than that of the vehicle-treated mice (Table 2, Supplementary Information), although both groups consumed more food compared to the db+/- mice during the treatment regime. The average weight of the liver (Fig. 7 D,E) and epididymal fat pads (Fig. 7 F,G) was the highest for db/db mice given vehicle control and least for the db+/- mice, while that for the compound treated db/db mice was intermediate. Hematoxylin and eosin staining of liver and adipose tissue(Fig. 7 H,I) showed signs of adipocyte hypertrophy and hepatocyte ballooning in vehicle-treated db/db mice, which were absent in the db+/- and LTK-14A treated db/db mice.

**Fig. 7:**
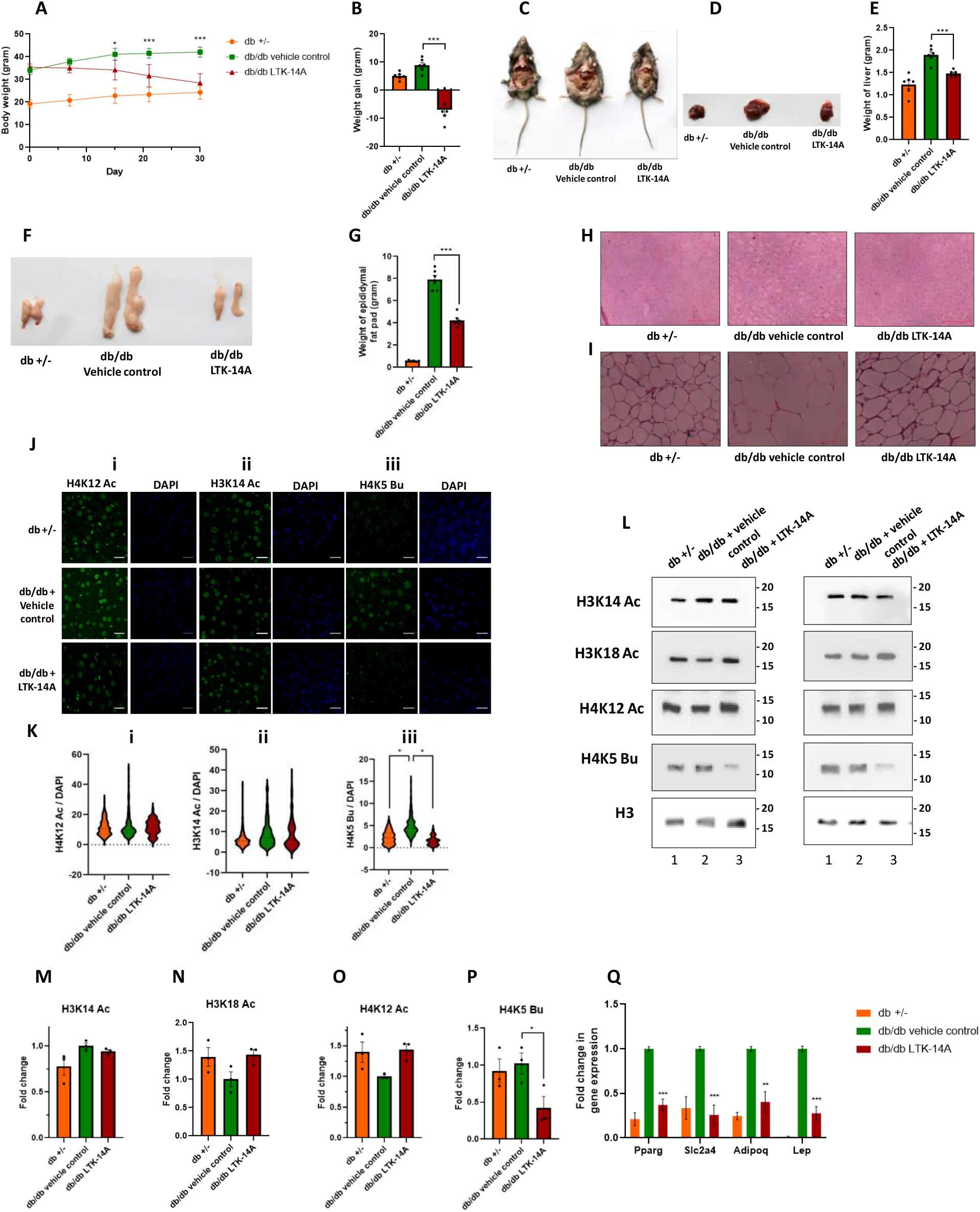
Oral gavaging of the butyrylation inhibitor reduces weight of genetically obese db/db mice: Trend of weight gaining (A) and average body weight gained after 30 days (B) of db +/- and db/db mice orally administered with either LTK-14A or vehicle solution; n=6 in each group. (C) Representative images of dissected bodies showing fat accumulation (panel iii) of the mice from three groups. Representative images of liver (D) from the mice of three groups and average weight of liver (E) depicted graphically; n= 6 in each group. Representative images of epididymal fat pads (F) from the mice of three groups and average weight of fat pads (G) depicted graphically; n= 6 in each group. Representative images of morphology of liver (H) and adipose tissue (I) from the mice of the three groups, as studied by hematoxylin and eosin staining. (J) Immunofluorescence microscopy of liver sections stained with antibodies against histone H4K12 acetylation (panel i), histone H3K14 acetylation (panel ii) and histone H4K5 butyrylation (panel iii). The fluorescence intensity of the modified histone marks was normalised against DAPI staining of whole nuclei for quantitation which is graphically depicted in (K) panel i, ii and iii ;n=4 in each group; one-way ANOVA with Dunnett’s multiple comparision: * P < 0.05, **P < 0.01, ***P < 0.001, ns: not significant. (L) Histone H3K14, H3K18, H4K12 acetylation and H4K5 butyrylation levels in epididymal fat pads of mice were estimated by immunoblotting. Histone H3 was used as loading control The fold change of quantified band intensity is graphically represented in panel M, N, O and P; n=3 in each group; one-way ANOVA with Dunnett’s multiple comparision: * P < 0.05, **P < 0.01, ***P < 0.001, ns: not significant. (Q) qRT-PCR with RNA isolated from epididymal fat pads to check the expression pattern of lipogenesis related pro-adipogenic genes upon LTK-14A treatment; n=3/4 in each group; one-way ANOVA with Dunnett’s multiple comparision: * P < 0.05, **P < 0.01, ***P < 0.001, ns: not significant

Similar to the high fat diet-induced obesity model, the more obese homozygous leptin receptor mutant mice (db/db) liver had a greater level of H4K5 butyrylation compared to leaner heterozygous mutant mice (db +/-) (Fig 7J panel iii, Fig. 7K panel iii). Moreover, LTK-14A treated db/db mice liver showed a decrease in H4K5 butyrylation compared to vehicle control-treated mice liver. In contrast, H4K12 (Fig. 7J panel i, Fig. 7K panel i) and H3K14 acetylation (Fig. 7J panel ii, Fig. 7K panel ii) were only partially upregulated in the db/db mice and did not get significantly reduced upon LTK-14A treatment. Immunoblotting with acid extracted histones from epididymal fat pads showed that H4K5 butyrylation was reduced in the adipose tissues of LTK-14A treated mice (Fig. 7 L,P) compared to vehicle control treated obese mice. H3K14, H3K18 and H4K12 acetylation were not significantly altered in any of the three groups (Fig. 7 L,M,N,O). Finally, RT-qPCR analysis showed that just like in cellular model of adipogenesis, LTK-14A treatment led to robust inhibition of important pro-adipogenic genes *Pparg*, *Lep*, *Slc2a4* and *Adipoq* in the fat pads of compound treated db/db mice (Fig. 7Q). Collectively these data establish that obesity can not only be prevented but also reverted by specific inhibition of H4K5 butyrylation, thereby highlighting the importance of this modification in the context of adipogenesis.

### LTK-14A is non-mutagenic and non-toxic in nature for biological applications

Based on the observations of obesity attenuating effects of LTK-14A in two different mice models, we performed some preclinical tests to ensure that the obesity arrest was not a result of any deleterious side effect of toxicity from the compound. Plate incorporation assay was carried out in four different nutritional auxotrophic mutants of Salmonella typhimurium (TA97a, TA98, TA100 & TA102) in the absence or presence of rat liver extract (S9 enzyme fraction). It was observed that the number of revertants obtained from LTK-14A was similar to that from vehicle control and far less compared to that from the positive control in all four strains, both in the absence and presence of liver extract (Sup. Fig 4 A-H). Furthermore, acute toxicity test was performed on rats for checking whether a single administration of excessively high dose of LTK-14A would alter the body weight and food/water consumption pattern of the rats. The body weight changes in the LTK-14A administered rats were similar to the vehicle control-treated ones (Table 6) with no gross morphological changes in any organ. Moreover, there was also no significant treatment-related effect on the food and water consumption of rats of both gender for all three doses compared to vehicle control (Table 6).

Collectively these data imply that LTK-14A does not have deleterious toxic effects on the test animals and could have a potential for downstream applications for tackling obesity-related complications.

## Discussion

Reports on the physiological relevance of acylation modifications are scanty. Breakthroughs have been made in establishing the importance of butyrylation in spermatogenesis (5), β-hydroxybutyrylation in starvation response (11) and crotonylation in inflammatory response, digestive system, cancer and HIV latency (7,12,13,14). Lysine propionylation has also been reported as an activation mark on histones that could be useful in various physiological processes (15). The list of these modifications seems to be increasing continuously, with the recent additions of histone serotonylation (16), lactylation (17) and dopaminylation (18). The fact that serotonin and dopamine are chemical neurotransmitters while lactate is derived from pyruvate, a glycolytic end product, and all these molecules bring about a modification of histones thereby mediating gene expression in several tissues, bear testimony to the hypothesis that the abundance levels of cellular metabolites in different physiological contexts can lead to modification of histones with a plethora of diverse chemical moieties that can selectively modulate gene expression patterns.

Our initial experiments in 3T3L1 adipogenesis indicated an important role of ACSS2 in the differentiation process by generating butyryl CoA for histone butyrylation. Inside the cells ACL (ATP citrate lyase) and ACSS2 have been implicated in the generation of nuclear acetyl CoA. Both the enzymes are known for regulating gene expression in different contexts by acetylating histones in specific genomic loci (19–21). Knockdown of *Acss2* had a much more pronounced effect on H4K5 butyrylation compared to H3K18 acetylation. This is probably due to the fact that ACSS2 is the sole/major producer of acyl CoA species while acetyl CoA can be generated by diverse means and the knockdown of a single acetyl CoA producer was not enough to have a robust effect on global acetylation level of histones. Similar observations have already been made for histone crotonylation in HeLa cells (7).

Due to redundancy in the functionality of p300 catalyzed butyrylation with acetylation, the significance of butyrylation in adipogenesis was demonstrated using a pharmacological approach. LTK-14A could selectively and specifically inhibit histone butyrylation and still inhibit adipogenesis by downregulating the expression of many pro-adipogenic genes. However amongst these genes *CEBPd* expression did not get affected by LTK-14A possibly because the expression of this gene is less prone to be regulated by histone butyrylation in its promoter. Indeed, our chromatin immunoprecipitation study indicated that in the initial stage of induction of differentiation in pre-adipocytes, there was no enrichment of H4K5 butyrylation in the promoter of *Cebpd*. This could explain why the butyrylation inhibitor LTK-14A treatment at the induction stage did not inhibit *Cebpd* expression, since there was no buyrylation mark enrichment for inhibition in its promoter to begin with. An increased butyrylation in its promoter at the terminal stage could be either through p300 catalysed modification or non-enzymatic means with greater availability of butyryl CoA, or both. Currently, we have not been able to verify this. However our preliminary observations with promoter-specific studies indicate that H4K5 butyrylation plays a role in regulating the expression of the critical transcription factor *Pparg*. Inhibition of butyrylation by LTK-14A repressed the expression of the master regulator which further led to the inhibition of downstream genes like *Lep*, whose expression is dependent on PPARγ. While cell line-based studies indicate that C/EBPδ regulates the expression of a cascade of transcription factors like C/EBPα and PPARγ, this fact has been questioned in knockout mice which have poorly differentiated adipose tissue but still express *Pparg* and *Cebpa* (22). This indicates that there could be other factors playing a redundant function like C/EBPδ in the initiation stage of adipogenesis. Moreover, an additional function of C/EBPδ may seem to be downstream of C/EBPα and PPARγ at the terminal stage of adipogenesis. It has been speculated that this could involve inducing the expression of ligands that activate PPARγ (23). Therefore, the increased butyrylation in the promoter of *Cebpd* on day 6 of adipogenesis might be responsible for its role in the terminal stage. The expression of *Cebpd* is independent of *Pparg* and hence repression of *Pparg* by butyrylation inhibition with LTK-14A did not repress *Cebpd*.

*Acss2*, which encodes the enzyme responsible for butyryl CoA biosynthesis was also downregulated by LTK-14A which might have further contributed to reduced butyrylation of histones during inhibition of adipogenesis. Amongst the different histone modification patterns investigated, H4K5 butyrylation was consistently affected by LTK-14A in different cell line and mice models, while the other site specific butyrylation marks were relatively unaltered, indicating that there may be a hierarchy in the importance of these modifications in the regulation of adipogenesis. While our work was in progress, a new report was published in which it was demonstrated that inhibition of fatty acid synthesis as well as oxidation leads to a reduction in H4K5 butyrylation while acetylation remains unaffected (24). This further reinforces our own hypothesis that fatty acid metabolism is an important source for certain acyl CoAs such as butyryl CoA, derived from the short chain fatty acid butyric acid. Another work published during this period revealed that HBO1, a member of the MYST family of acetyltransferase, also possesses the ability to catalyse acylation reactions including butyrylation (25). Mass spectrometry studies of histone modifications altered by knockdown of HBO1 in HeLa cells showed that H4K5 butyrylation does not get affected by depletion of HBO1 indicating that it is a p300 specific mark. LTK-14, the monomethoxy derivative of isogarcinol was found to be a specific inhibitor of p300 acetyltransferase activity (8) without affecting PCAF. Since its disubstituted counterpart LTK-14A was found to be relatively inert with respect to LTK-14, it seems plausible that LTK-14A would be unlikely to affect other epigenetic modifications besides p300 catalysed acylation.

Global histone acetylation has previously been reported to remain unchanged in liver and adipose tissues of mice even after high fat diet consumption (26), which was consistent with the observations we made in both mice models studied. Since acetyl CoA is highly abundant, fluctuations in its levels did not lead to major changes in histone acetylation patterns. Because the usual butyryl CoA levels are far less than their corresponding Michaelis Menten constant for p300, an increase in butyryl CoA levels due to excess lipid consumption through high fat diet or *de novo* lipid synthesis through excess food consumption, resulted in a concomitant increase in butyrylation by p300 which served as the nutritional sensor for butyryl CoA. Enhanced H4K5 butyrylation was not observed in mature adipose tissues of obese mice possibly due to the predominance of adipocyte population in the tissues of mice from all the groups (27).

## Conclusion

In the 21st century, obesity has become a major epidemic leading to several other disorders such as type 2 diabetes and arteriosclerosis. Histone acetylation inhibition could be regarded as a possible method for attenuating adipogenesis and obesity. But acetylation is a widespread phenomenon that is required in several other physiological processes besides adipogenesis. Hence, treatment with an acetyltransferase inhibitor at a particular range of concentration could have undesirable side effects and toxicity. In contrast, butyrylation is a rarer event, repressing which may be less harmful to other processes. Indeed, our preclinical tests are indicative of the fact that LTK-14A may be suitable for development as a possible anti-obesity drug due to its negligible toxicity side effects. The present finding thus establishes the importance of histone butyrylation in adipogenesis, making it an attractive target for pharmacological inhibition of adipogenesis and thereby controlling obesity (Graphical abstract).

## Supplementary Section

### Supplementary Materials and Methods

#### siRNA experiments

ON-TARGETplus siRNA SMART pools against mouse Acss2 (catalog no.L-065412-01-0005) was purchased from Dharmacon (Dharmacon, Lafayette, CO, USA). 3T3L1 cells were transfected with Acss2 siRNA at 25 nM concentration, using RNAiMAX (catalog no. 13778-150, Thermo Fisher Scientific) according to manufacturer’s protocols. Transfection was carried out for 48 hours after which cells were induced for differentiation for 6 days followed by RNA isolation and histone extraction.

#### Molecular docking analysis

Glide module of Schrodinger suite was used to dock LTK-14 and LTK-14A to p300 catalytic domain (PDB: 3BIY). Chimera was used as visualizing software. The protein was prepared using the wizard tool in Maestro version 10.2. The crystal structure of LTK-14A (CCDC 1969185) and LTK-14 (CCDC 645420) were used as the most optimized and energy minimized ligand structures. The receptor grid for p300 catalytic domain was generated by specifying the binding (active) site residues using Receptor grid generation in the Schrodinger Glide application. Once the receptor grid was generated, the ligands LTK-14 and LTK-14A were docked to the p300 catalytic domain using Glide version 5.8 molecular docking.

#### Untargeted analysis of intracellular metabolites by Ultra-performance liquid chromatography coupled with time-of-flight mass spectrometer (Q-TOF LC/MS)

Profiling of intracellular metabolites was performed on an agilent 1290 Infinity LC system coupled to Agilent 6545 Accurate-Mass Quadrupole Time-of-Flight (QTOF) with Agilent Jet Stream Thermal Gradient Technology. The UPLC system was assembled with a Diode array detector (DAD) and autosampler. The Chromatographic separation was achieved on Agilent ZORBAX SB-C18 column (2.1 × 100 mm, 1.8 µm) as stationary phase. The mobile phase consisted of a linear gradient of 100 mM ammonium formate (A) and Acetonitrile (B): 0–10.0 min, 30-80% B (v/v); 10.0–15.0 min, 80–100% B (v/v); 15.0–20.0 min, 100% B (v/v); 20.0–21.0 min, 100–30% B (v/v); 21.0–25.0 min, 30% B. The sample was dissolved in 1 ml methanol (LCMS Grade), centrifuged and supernatant was taken for UPLC-QTOF-MS analysis. The column was reconditioned for 5 minutes prior to the next injection. The flow rate was 0.5 ml/min, and the injected volume was 20 µL. The UPLC was connected to the MS analysis was performed on an Agilent 6545 Accurate-Mass Q-TOF/MS system with an electrospray ionization (ESI) source. Considering the MS conditions, positive ion mode was used to obtain high-resolution mass spectra. The ESI source parameters were: drying gas (N2) flow, 8 L/min; drying gas temperature, 200°C. Other parameters were set as nebuliser gas, 35 psig; capillary voltage, 3000 V; skimmer voltage, 65 V; nozzle voltage 1000 V and fragmentor voltage 150 V. The data acquisition on the LC-QTOF was performed using Agilent Mass Hunter Acquisition software (Agilent Technologies, Santa Clara, CA, USA). The data were deconvoluted into individual chemical peaks with Agilent MassHunter Qualitative Analysis (MassHunterQual, Agilent Technologies).

#### Site directed mutagenesis

Point mutation was introduced into the p300 catalytic domain sequence cloned within pET-28b plasmid by using the Quikchange site directed mutagenesis kit (Agilent). Briefly, the template was mixed with Pfuturbo DNA polymerase, designed forward and reverse primers and dNTPs for PCR (Initial denaturation for 30 seconds at 95°C; Cycles: 95°C 30’’, 55°C for 60’’, 68°C for 7’; final elongation at 68°C for 5’.) The mixture was then incubated with DpnI enzyme for 2.5 hours for removal of the parent wild type sequence followed by transformation of competent XL10Gold cells and kanamycin selection of positive transformants. The inserted mutation was verified by sequencing and alignment with the wild type sequence using ClustalOmega software. The sequences of the primers used are as follows:

Forward primer-5’-CCTCACTTGGTGGAGCTGCCCAAAATATGCCCTGTTGTG – 3’

Reverse primer-5’-CACAACAGGGCATATTTGGGCAGCTCCACCAAGTGAGG -3’

#### Purification of recombinant full-length p300 and p300 catalytic domain

Polyhistidine tagged recombinant p300 was purified from Sf21 insect cells by Ni-NTA based affinity chromatography. These cells were infected by the baculovirus containing p300 for 72 hours. Once the cells showed signs of infection, they were scraped off and centrifuged to separate the spent media. The cell pellets were resuspended in ice cold homogenisation buffer (20 mM Tris, pH 7.5; 20 mM imidazole, 10% glycerol, 0.2 mM EDTA, 300 mM KCl, 0.1%NP-40, 2 mM PMSF, 2 mM βmercaptoethanol) and then homogenised using a Dounce homogeniser (5 cycles, 6 strokes per cycle with 5 minutes interval between each cycle). The cell debris was separated by centrifugation and then the lysate was incubated with pre-equilibriated Ni-NTA slurry for 2.5 hours at 4°C on an end-to-end rotor. Then the beads were washed with wash buffer (20 mM Tris, pH 7.5; 40 mM imidazole) nine times to remove unwanted contaminants. Finally p300 was eluted using elution buffer (20 mM Tris, pH 7.5; 250 mM imidazole).

Polyhistidine tagged p300 catalytic domain cloned in pET28b plasmid and Sirt2 were used for co-transforming BL21(DE3) cells. At first, 50 ml primary culture was prepared in Luria broth containing kanamycin (50 µg/ml) and ampicillin (100 µg/ml) (HIMEDIA) and then it was scaled up 10 times to prepare secondary culture. Incubation was done at 37⁰C, 180 rpm till O.D. reached 0.37 followed by induction with 0.2 mM IPTG and a further incubation at 30⁰C for 4 hours. Cells were pelleted at 6000 rpm, 4⁰C for 10 minutes and stored at −80⁰C. Resuspension and homogenisation of the pellet was carried out in homogenisation buffer (20 mM Tris, pH 7.5; 20 mM imidazole, 10% glycerol, 0.2 mM EDTA, 300 mM KCl, 0.1%NP-40, 2 mM PMSF, 2 mM βmercaptoethanol). 3 cycles of sonication were done for 3 minutes at 35% amplitude with 5 minutes interval between each cycle. The cell debris were separated by pelleting at 12000 rpm for 30 minutes at 4⁰C. The supernatant was equilibriated with Ni-NTA beads slurry (Novagen) in an end-to-end rotor for 3 hours at 4 C. Eight washes were given with wash buffer (20 mM Tris, pH 7.5; 40 mM imidazole). Elution was carried out in batches with elution buffer (20 mM Tris, pH 7.5; 250 mM imidazole).

The enzymatic activity of the proteins was estimated by filter binding assay.

#### *In vitro* filter binding assay

Different dilutions of purified enzyme were incubated in reaction buffer (50 mM Tris-HCl pH 7.5, 1 mM DTT, 10 % Glycerol, 2 mM PMSF) with 10 mM NaBu, tritiated acetyl CoA (50 µCi) (PerkinElmer, Part No: NET290050UC) and Xenopus histone H3 (500 ng) for 30 minutes at 30⁰C. The reaction mixture was then spotted on P81 filter paper and the paper was allowed to dry at room temperature. The paper was washed in wash buffer (50 mM sodium bicarbonate and 50 mM sodium carbonate), dried at 90ᴼC and immersed in enhancing solution (2.5% PPO and 0.25% POPOP in toluene) for taking radioactive counts using liquid scintillation counter (Perkin Elmer, MicroBeta2 system).

#### *In vitro* acylation (acetylation/butyrylation) assay

Acylation reactions were performed in reaction buffer (25 mM Tris-HCl pH 7.5, 100 mM NaCl, 0.1 mM EDTA, 1 mM DTT, 10 % Glycerol, 1x PIC) with 100 ng/mL TSA, and 50 μM Butyryl-CoA/Acetyl CoA. Xenopus histone H3 (1 µg) was used as the substrate and full length p300 or p300 catalytic domain (10,000 cpm activity) was used as the enzyme. Reactions were incubated in presence or absence of butyrylation inhibitor for 10 minutes at 30 °C, followed by initiation of the acetylation/butyrylation reaction by the addition of acetyl CoA/butyryl CoA. After a further 10 minutes, reactions were stopped by addition of Laemmli buffer and samples were used for immunoblotting.

#### Transcriptomics analysis by RNA-seq

RNA integrity was checked by Agilent Bioanalyzer 2100, only samples with clean rRNA peaks were used. Libraries for RNA-seq were prepared according to KAPA Stranded RNA-Seq Kit with RiboErase (KAPA Biosystems, Wilmington, MA) system. Final library quality and quantity were analyzed by Agilent Bioanalyzer 2100 and Life Technologies Qubit3.0 Fluorometer, respectively. 150 bp Paired-end sequencing was performed on Illumina HiSeq 4000 (Illumnia Inc., San Diego, CA).

House mouse (Mus musculus, strain C57BL/6J) genome (mm10) was downloaded from GENCODE and indexed using Bowtie2-build with default parameters. Adapter removal was done using Trim Galore (v 0.4.4) and each of the raw Fastq files were passed through a quality check using FastQC. PCR duplicates were removed using the Samtools 1.3.1 with the help of ‘rmdup’ option. Each of the raw files was then aligned to mm10 genome assembly using TopHat2 with default parameters for paired-end sequencing as described in (28). After aligning, quantification of transcripts was performed using Cufflinks and then Cuffmerge was used to create merged transcriptome annotation. Finally differentially expressed (DE) genes were identified using Cuffdiff. The threshold for DE genes was log2 (fold change) >1.5 for up regulated genes and log2 (fold change) <1.5 for down regulated genes with p-value <0.05.

##### GO enrichment analysis

Gene Ontology (GO) analysis was performed in PANTHER (29). Significant enrichment test was performed with the set of differentially expressed genes in PANTHER and Bonferroni correction method was applied to get the best result of significantly enriched biological processes.

##### Fisher’s exact test

Fisher’s exact test was performed in PANTHER Gene Ontology (GO) where p-value significance was calculated based on the ratio of obtained number of genes to the expected number of genes (O/E) considering the total number of genes for the respective pathway in Mus musculus with a FDR of <0.05.

##### Heatmap and clustering of genes

Unsupervised hierarchical clustering method was performed using Cluster 3.0 (30) with Pearson Correlation and average linkage rule. Gene expression data (FPKM of all samples) was taken and log2 transformed. Low expressed (FPKM<0.05) and invariant genes were removed. Then genes were centered and clustering was performed based on differential expression pattern of genes and fold change. Finally, the heatmap was visualized in Java TreeView 3.0.

##### Biological analysis of differentially expressed transcripts and pathway regulatory network modeling

Statistically significant differentially expressed transcripts were subjected to GO and Pathway enrichment using DAVID tool. Only those GO and pathways with a FDR score of <=0.05 was considered for further downstream analysis. Key biologically dysregulated GO and Pathways along with the differentially expressed genes was provided as an input to Pathreg algorithm from Theomics International Pvt Ltd, Bangalore, India for gene regulatory network modeling. The result (nodes and edges) of the Pathreg algorithm was provided as an input to Cytoscape v2.8.2 to identify key nodes and edges that could be representative of the gene regulatory changes upon treatment.

#### Quantitative real time PCR

Total RNA was isolated from the 3T3L1 cells and epididymal fat pads by phenol-chloroform method. In case of LTK-14A treatment, 3T3L1 cells were allowed to differentiate for 7 days following the standard adipogenesis protocol, in presence of the compound at 25 µM concentration. The control experiments were 3T3L1 cells differentiated with or without DMSO as solvent control and differentiation was done only with the addition of only the cocktail of chemicals for adipogenesis. RNA was isolated from the cells after the period of 7 days in all three cases. For isolation of RNA from mice fat pads, the adiose tissue samples were lysed in Trizol using a mechanical homogenizer followed by phenol-chloroform method of extraction. The isolated RNA samples were treated with 2 units/µL DNase I to remove any residual DNA followed by ethanol precipitation at −80ᵒC. 2μg RNA was used for cDNA synthesis using MMLV reverse transcriptase (Sigma) in presence of 0.5 mM dNTP and 3.5 µM oligo dT (Sigma). RT-qPCR amplification was done in kappa SYBR green reagent (Biosystems) with the help of Real time PCR instrument (Applied Biosystems by Life Technologies). For estimation of relative fold change in RNA expression, the following calculation was done: Relative fold change = 2-ΔΔCt, where ΔΔCt=ΔCt value of sample – ΔCt value of control and ΔCt= Ct value of target – Ct value of internal control actin for individual sample. The sequences of primers are provided in Table 4.

**Table 3:**
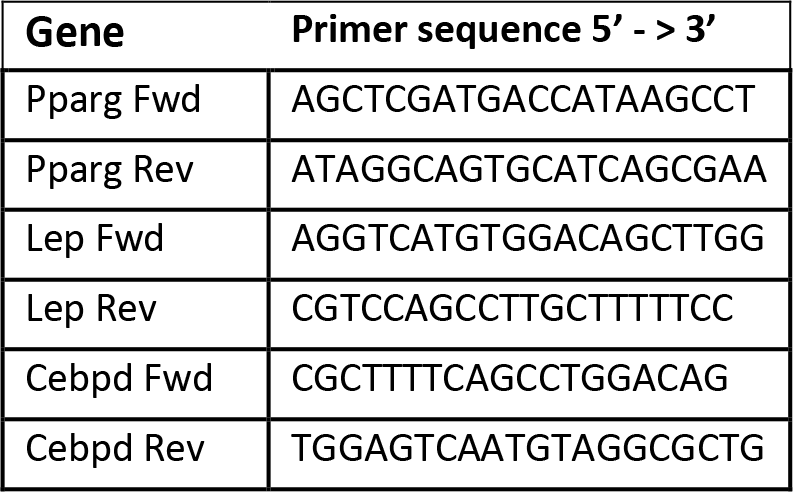
List of primers used in ChIP-qRTPCR

**Table 4:**
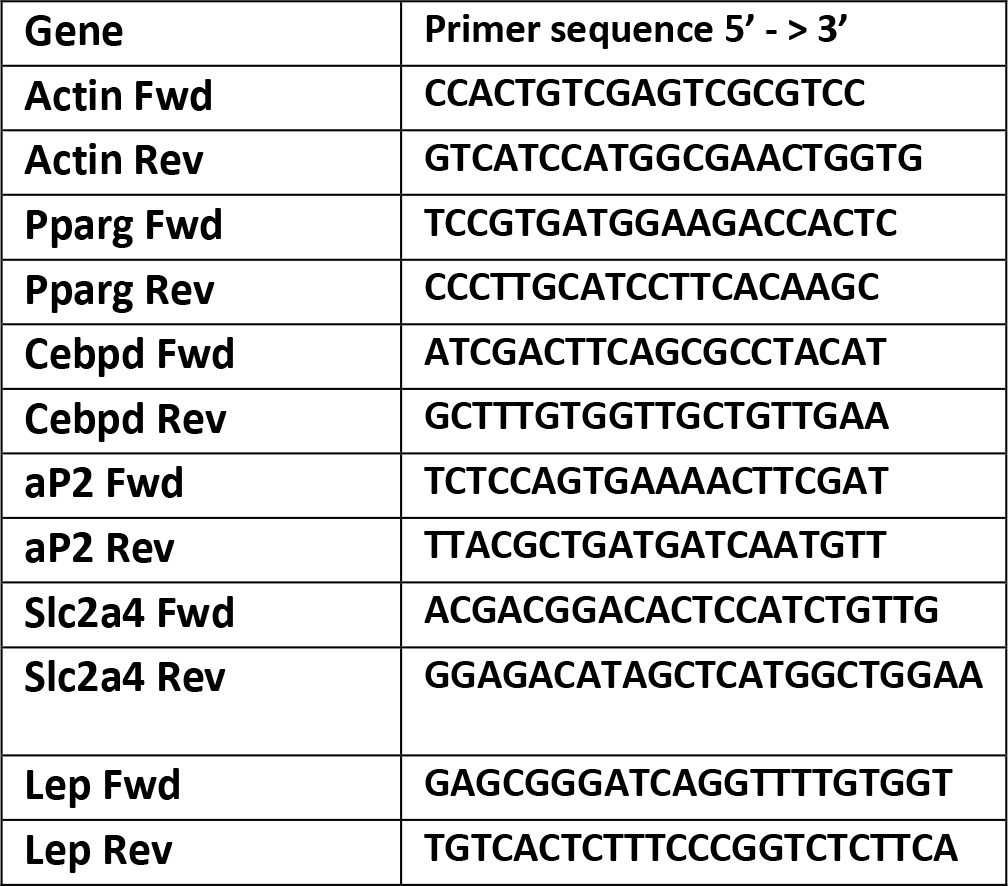
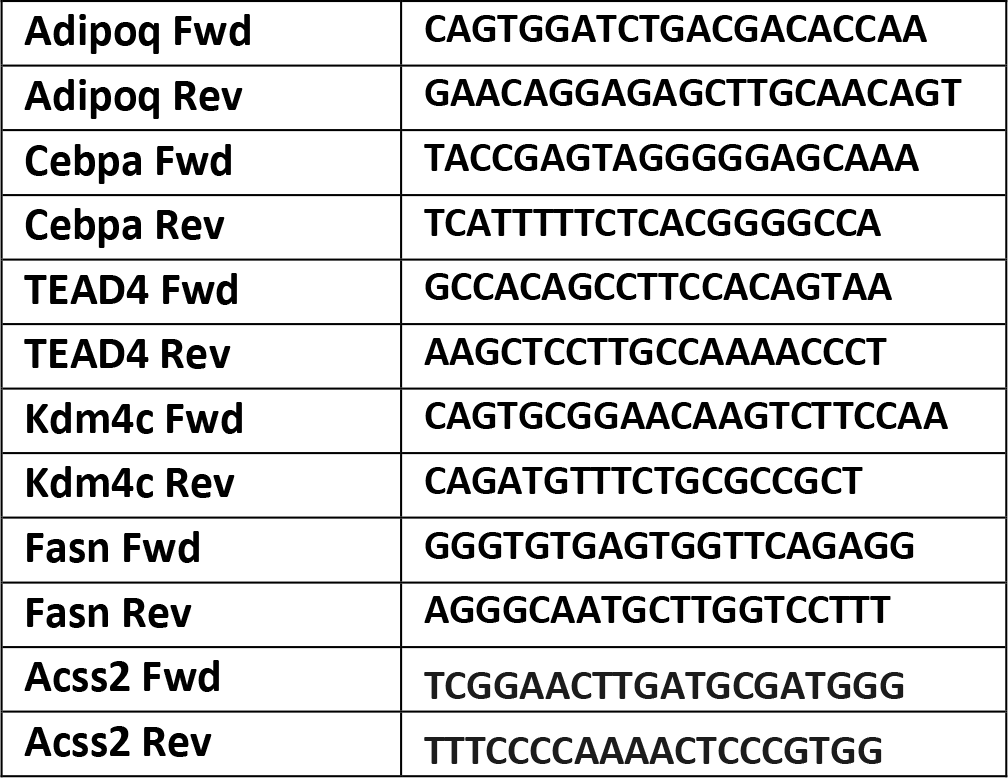
List of primers used in gene expression analysis by qRTPCR

#### Hematoxylin and eosin staining of liver and adipose tissues

Staining was performed on paraffin embedded sections of liver and epididymal fat pads having the thickness of 5 µm and 10 µm respectively. Briefly, the sections were deparaffinized in xylene, followed by immersion in absolute alcohol, 90% ethanol, 70% ethanol, 50% ethanol and water before staining with hematoxylin for 5-10 minutes. The sections were immersed in above mentioned solutions in reverse order before counter-staining in eosin for 15 seconds. Then the samples were dipped in absolute ethanol and xylene before mounting with DPX.

#### MTT assay

12500 cells/ml were seeded in presence/absence of LTK-14A in 96-well plate and at different intervals of time (2 days, 4 days, 6 days), 1/10th volume of 5mg/ml MTT solution was added to it followed by incubation at 37 °C for 3 hours. The formed crystals were dissolved in DMSO and spectrometric estimation of colour intensity was done at 570 nm.

#### Evaluation of mutagenecity by plate incorporation assay

Five doses of LTK-14A (10.0□g, 33.0□g, 100.0□g, 333.0□g, and 1000.0□g/plage) were selected to check for its possible mutagenecity. Seven groups decided as Group 1-Control, Group 2 Positive Control and 5 doses of LTK-14A (Group 3 to Group 7: 10.0□g, 33.0□g, 100.0□g, 333.0□g and 1000.0□g/plate) were tested. Four tester strains of Salmonella typhimurium (TE 97a, TA 98, TA 100, TA102) were used and their respective positive controls were used in absence of S9 fraction (Table 5); 2-aminoanthracene was used as the positive control in presence of S9 fraction (Table 5). The tester strains were grown overnight in Luria Broth antibiotic selection medium at 37°C. LTK-14A prepared in DMSO was mixed with the bacterial culture, 0.5mM Histidine-Biotin and top agar in a culture tube and the content was overlaid on to minimal glucose plate. In case of plate incorporation assay with S9 mix, LTK-14A was incubated with S9 mix in cold condition for 15 to 20 minutes followed by the same procedure as the plate incorporation assay without S9. The plates were incubated in an inverted position in 37°C incubators for 48 hours following which the number of revertants was counted using automated colony counter.

**Table 5:** List of mutagenic agents used as positive control in Ames test

| **Tester Strains** | **Without S9** | **With S9** |
| --- | --- | --- |
| **TA-97a** | 9-aminoacridine<br>(50µg/plate) | 2-aminoanthracene<br>(5µg/plate) |
| **TA-98** | N-nitro-o-phenylene<br>diamine<br>(2.5µg/plate) | 2-aminoanthracene<br>(5µg/plate) |
| **TA-100** | Sodiumazide<br>(5µg/plate) | 2-aminoanthracene<br>(5µg/plate) |
| **TA-102** | Mitomycin C<br>(0.5µg/plate) | 2-aminoanthracene<br>(5µg/plate) |

**Table 6:**
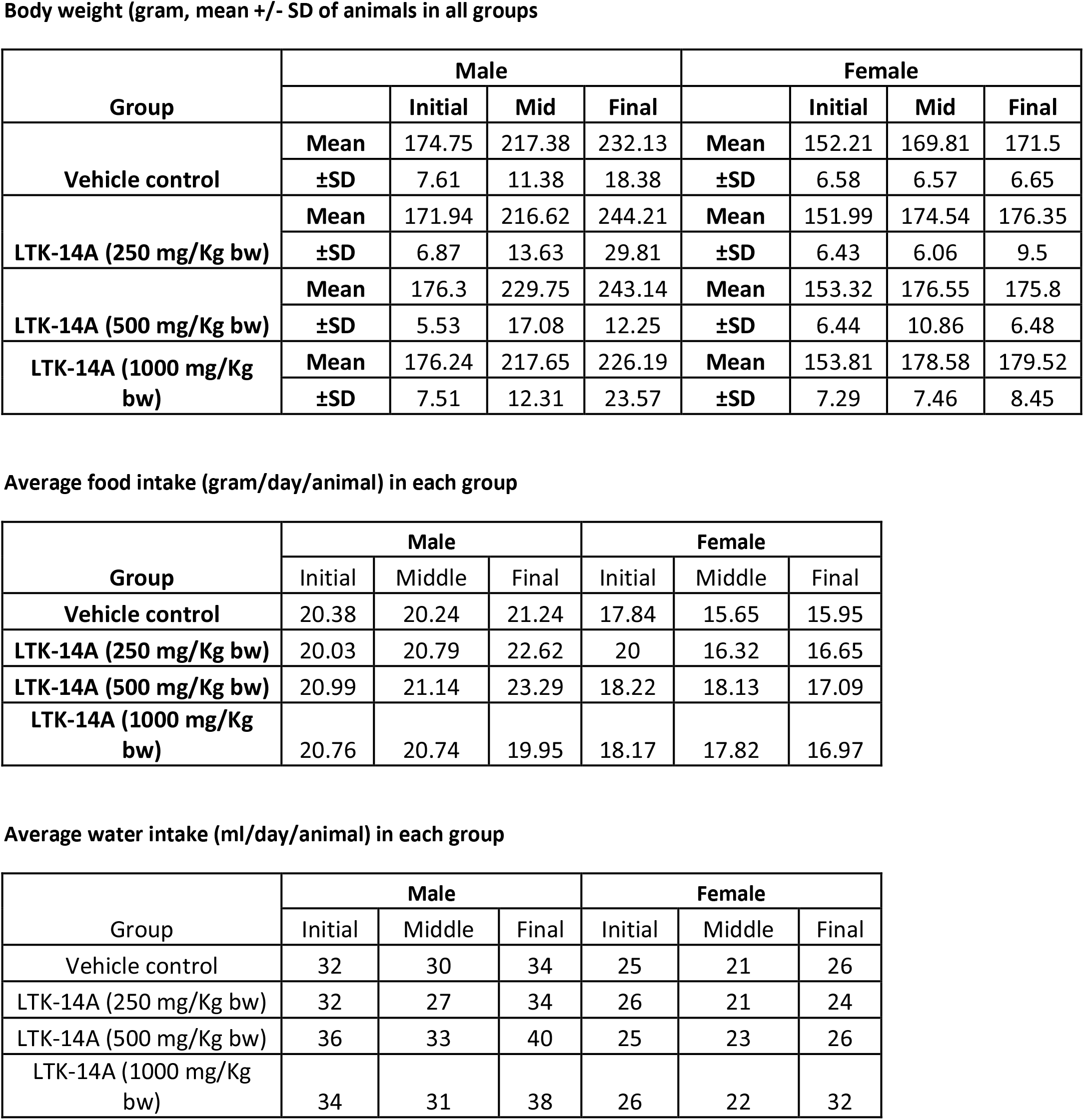
Details of single dose toxicity test

**Body weight (gram, mean +/- SD of animals in all groups**
| Group | Male |  |  |  | Female |  |  |  |
| --- | --- | --- | --- | --- | --- | --- | --- | --- |
|  |  | Initial | Mid | Final |  | Initial | Mid | Final |
| Vehicle control | Mean | 174.75 | 217.38 | 232.13 | Mean | 152.21 | 169.81 | 171.5 |
|  | ±SD | 7.61 | 11.38 | 18.38 | ±SD | 6.58 | 6.57 | 6.65 |
| LTK-14A (250 mg/Kg bw) | Mean | 171.94 | 216.62 | 244.21 | Mean | 151.99 | 174.54 | 176.35 |
|  | ±SD | 6.87 | 13.63 | 29.81 | ±SD | 6.43 | 6.06 | 9.5 |
| LTK-14A (500 mg/Kg bw) | Mean | 176.3 | 229.75 | 243.14 | Mean | 153.32 | 176.55 | 175.8 |
|  | ±SD | 5.53 | 17.08 | 12.25 | ±SD | 6.44 | 10.86 | 6.48 |
| LTK-14A (1000 mg/Kg bw) | Mean | 176.24 | 217.65 | 226.19 | Mean | 153.81 | 178.58 | 179.52 |
|  | ±SD | 7.51 | 12.31 | 23.57 | ±SD | 7.29 | 7.46 | 8.45 |

**Average food intake (gram/day/animal) in each group**
| Group | Male |  |  | Female |  |  |
| --- | --- | --- | --- | --- | --- | --- |
|  | Initial | Middle | Final | Initial | Middle | Final |
| Vehicle control | 20.38 | 20.24 | 21.24 | 17.84 | 15.65 | 15.95 |
| LTK-14A (250 mg/Kg bw) | 20.03 | 20.79 | 22.62 | 20 | 16.32 | 16.65 |
| LTK-14A (500 mg/Kg bw) | 20.99 | 21.14 | 23.29 | 18.22 | 18.13 | 17.09 |
| LTK-14A (1000 mg/Kg bw) | 20.76 | 20.74 | 19.95 | 18.17 | 17.82 | 16.97 |

**Average water intake (ml/day/animal) in each group**
| Group | Male |  |  | Female |  |  |
| --- | --- | --- | --- | --- | --- | --- |
|  | Initial | Middle | Final | Initial | Middle | Final |
| Vehicle control | 32 | 30 | 34 | 25 | 21 | 26 |
| LTK-14A (250 mg/Kg bw) | 32 | 27 | 34 | 26 | 21 | 24 |
| LTK-14A (500 mg/Kg bw) | 36 | 33 | 40 | 25 | 23 | 26 |
| LTK-14A (1000 mg/Kg bw) | 34 | 31 | 38 | 26 | 22 | 32 |

#### Single dose toxicity study

The single dose toxicity study on LTK-14A was conducted in accordance with Good Laboratory Practice Principles as published by the OECD in 1998, in accordance with the Drugs and Cosmetics Rule (New Drugs and Clinical trials, 2019/ DCGI). LTK-14A (formulated in vehicle i.e. 0.5% sodium salt of carboxymethyl cellulose) was administered to Sprague Dawley rats once by oral route on the day of scheduled treatment. The three treatment groups were 250 mg, 500 mg, and 1000 mg/kg body weight, while vehicle control was administered to control group. The rat equivalent doses were calculated from the dosage used in mice experiments following FDA guidelines for drug administration. The doses finally administered were 10, 20 and 40 times the corresponding mice equivalent dose. Food, but not water, was removed thus letting the animals fast over night prior to dosing. After administration of LTK-14A, food was withheld for further 2-3 hours and the animals were observed at 30 minutes, 1 hour, 2 hours and 4 hours from the time of dosing. Animals were subsequently observed once daily for any signs of toxicity, mortality, or any other observations including conditions of skin, hair coat, eyes, mucous membrane, movement, activity, tremors, convulsions, salivation, diarrhea, posture, piloerection and lethargy. Food intake, water intake and body weight of animals was recorded weekly. Terminal sacrifice of surviving animals was performed after 14 days using CO2 euthanasia technique following which animals were necropsied. Animals were fasted over night before necropsy. Gross observations were recorded.

---

**Sup. Fig.1:**
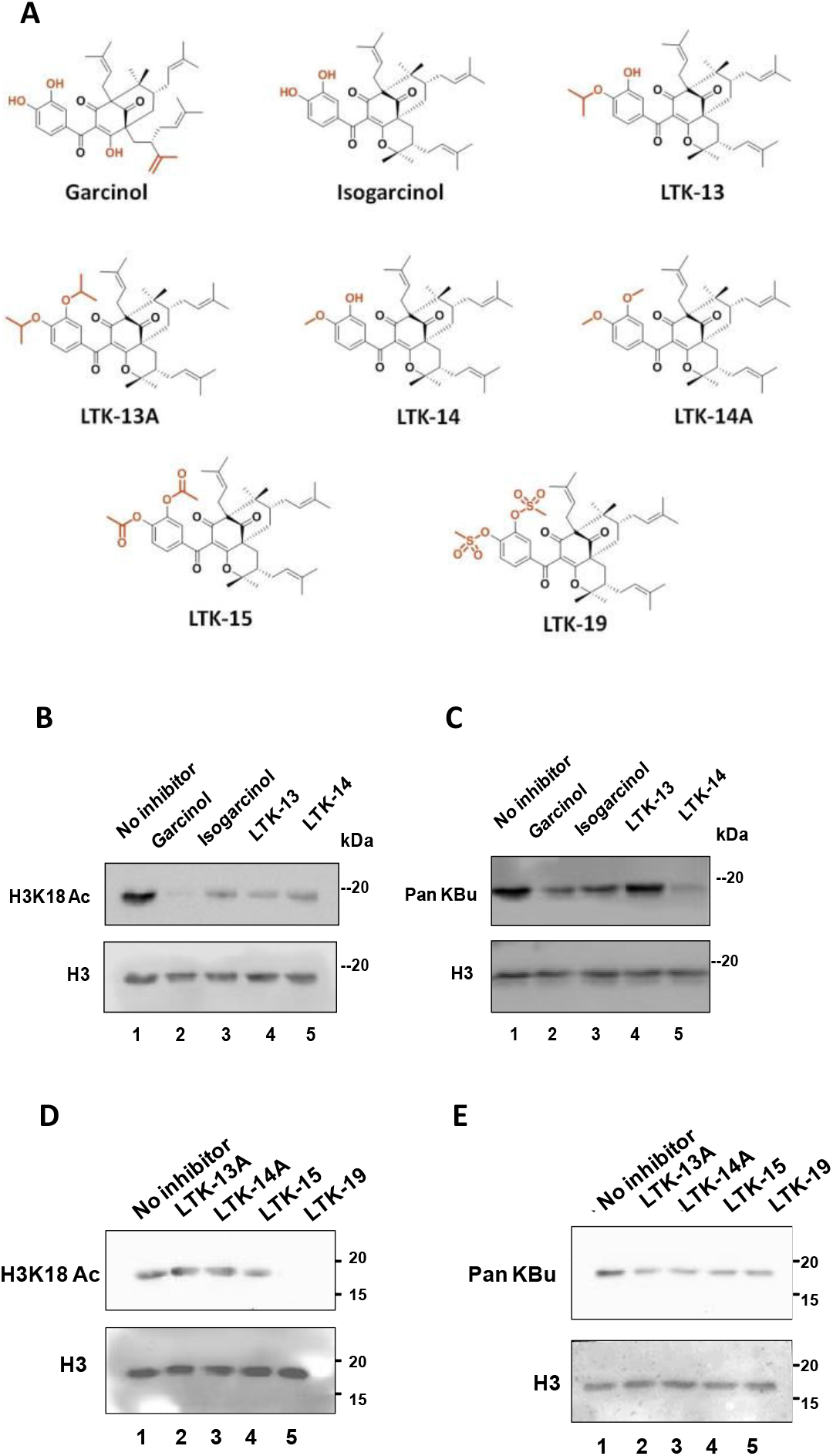
(A) Structures of garcinol and its cyclised and substituted derivatives. The site of internal cyclisation and mono/di-substitutions are shown in brown colour. Initial screening of the different monno-substituted derivatives for their differential effect on p300 catalysed acetylation (B) and butyrylation (C) of histone H3 at a concentration of 10 μM. Immunoblotting with antibodies against acetylated H3K18 (B) and pan-butyryl lysine (C). Similar screening was performed with di-substituted derivatives for their effect on p300 catalysed acetylation (D) and (E) of histone H3 at a concentration of 25 μM.

**Sup. Fig.2:**
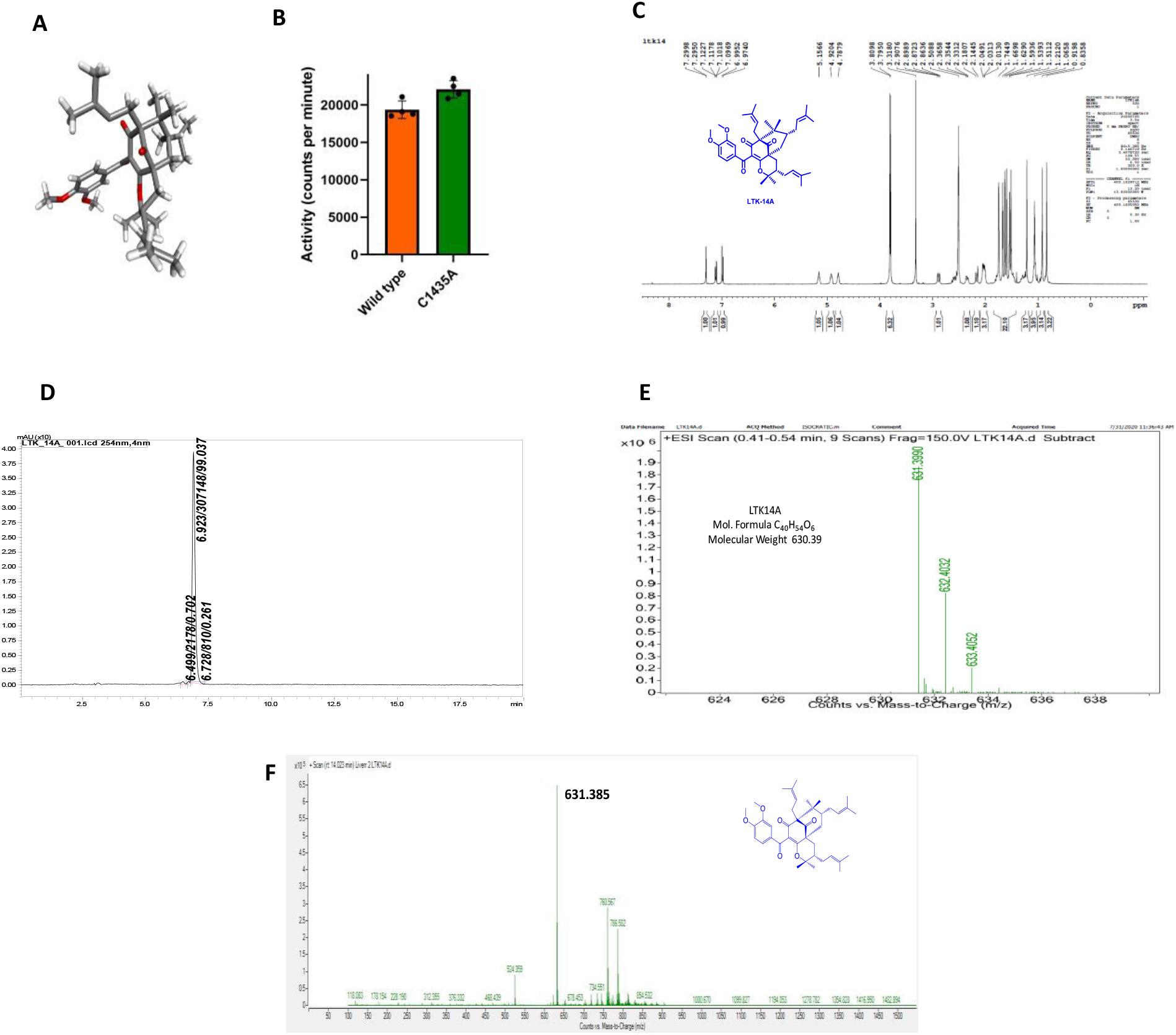
(A) Crystal structure of LTK-14A (CCDC: 1969185) (B) Bar graphs depicting the acetyltransferase activity of wild type an mutant C1438A p300 catalytic domain as meeasured by filter binding assay. Error bars denote mean +/- SEM of four technical replicates; two-tailed unpaired Student’s t-test: * P < 0.05, **P < 0.01, ***P < 0.001, ns: not significant. (C) Proton NMR, (D) HR-MS spectra and (D) HPLC chromatogram of LTK-14A. (F) The presence of LTK-14A in the physiological system upon administering the compound was verified by LC-ESI-MS analysis of the metabolite extracts of liver.

**Sup. Fig. 3:**
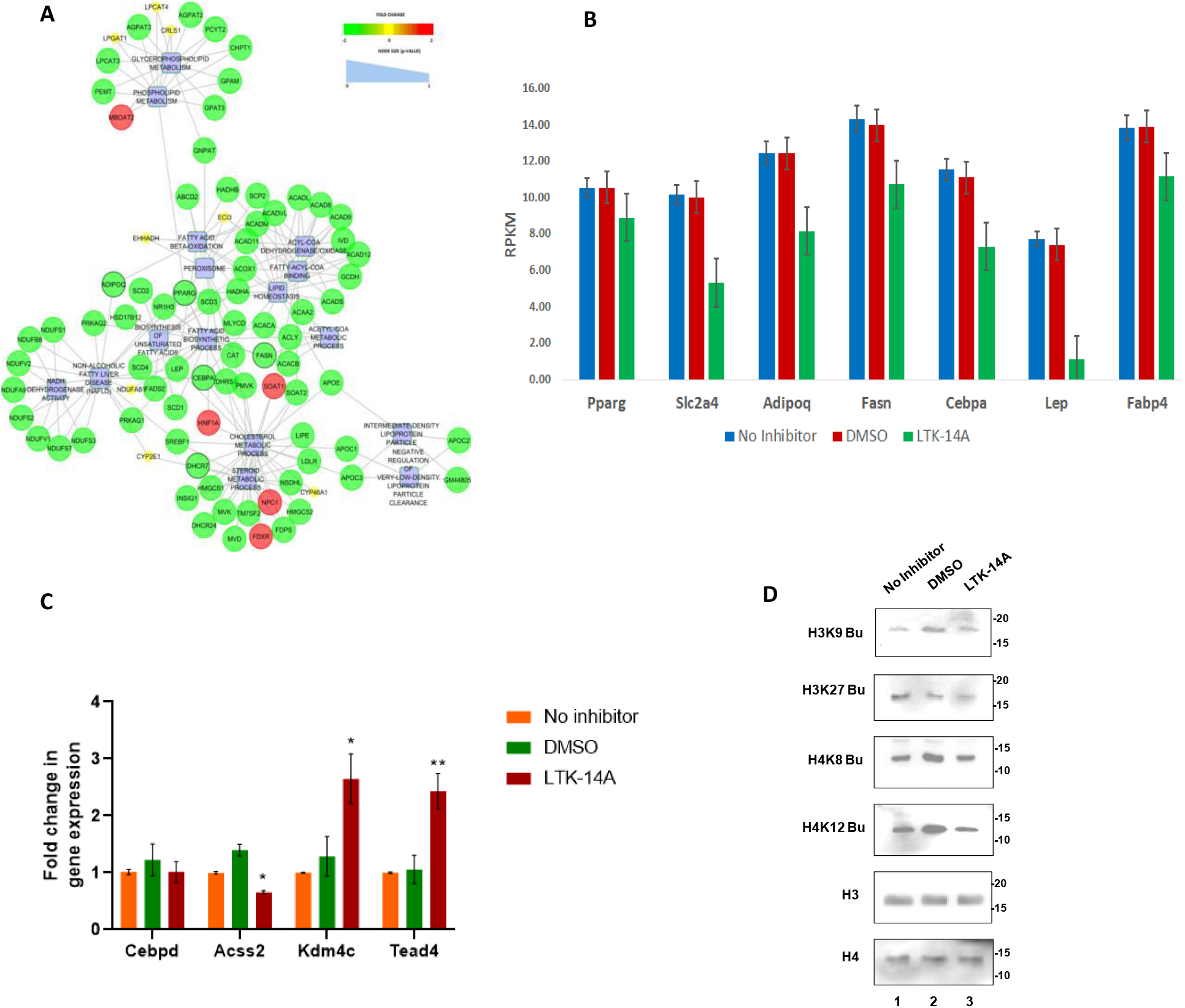
(A) Molecular pathway analysis for the expression status of major lipid metabolism related genes and their interrelationship with the pathways that were affected in LTK-14A versus untreated conditions. (B) Pro-adipogenic genes that could be putative targets for LTK-14A were identified and RPKM values for these genes have been depicted graphically.(C) RT-qPCR analysis was performed to validate the putative target genes of LTK-14A. Error bars denote mean +/- SEM of three biological replicates. (D) Histone butyrylation levels in 3T3L1 cells that were treated with LTK-14A were compared with that in DMSO treated and untreated conditions by immunoblotting with antibodies against butyrylated histone H3K9, H3K27, H4K8 and H4K12. Histones H3 and H4 were used as loading controls.

**Sup. Fig.4:**
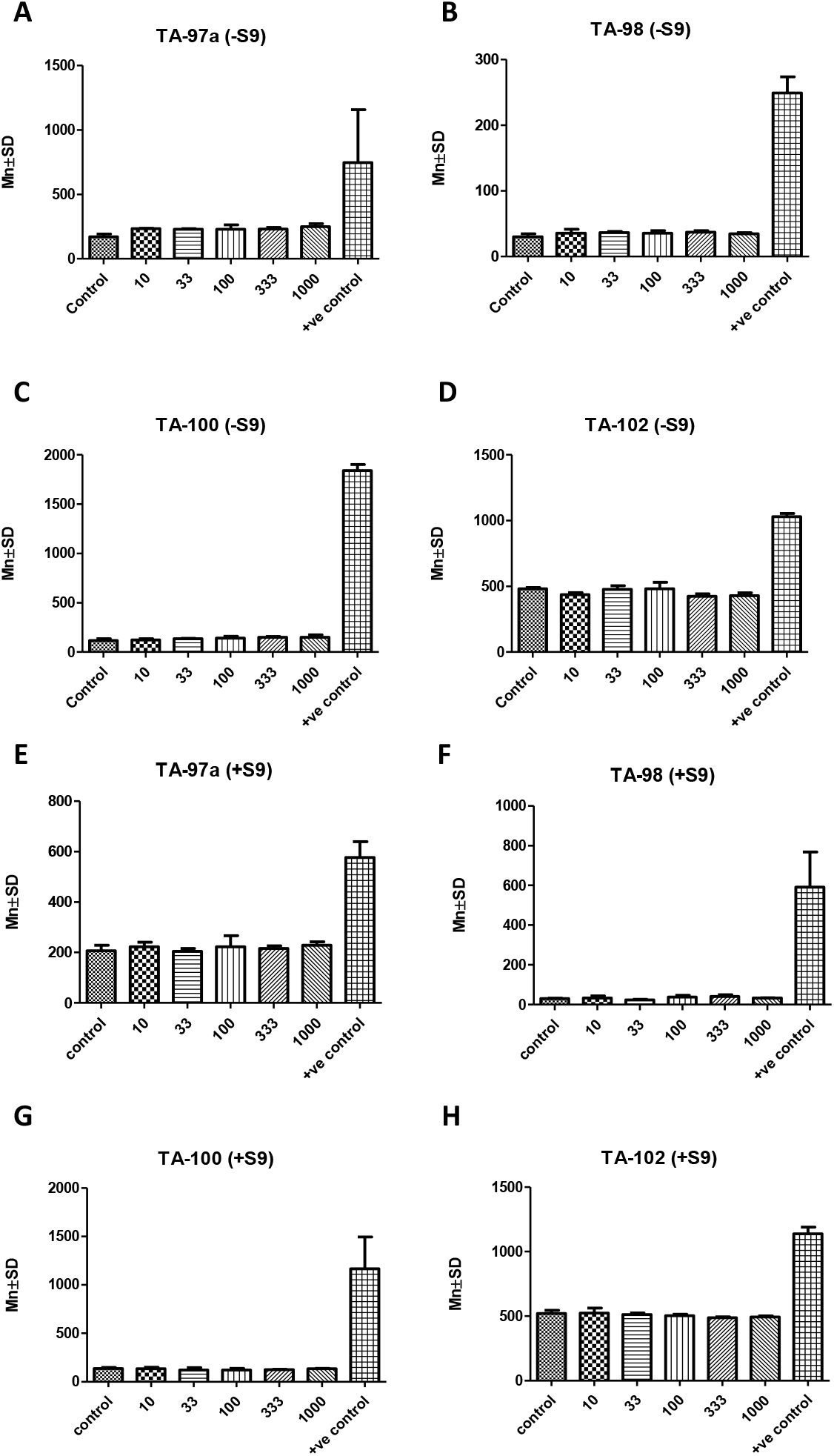
Evaluation of mutagenecity of LTK-14A at different doses was tested by performing plate incorporation assay with four different tester strains of Salmonella typhimurium (TE 97a, TA 98, TA 100, TA102). Panel (A) to (D) – tests without S9 fraction. Panel (E) to (H) – tests with S9 fractions. Error bars denote mean +/- SEM of three biological replicates; One-way ANOVA followed by Neman-Keuls test.

